# Chromosome-scale assemblies reveal the structural evolution of African cichlid genomes

**DOI:** 10.1101/383992

**Authors:** Matthew A. Conte, Rajesh Joshi, Emily C. Moore, Sri Pratima Nandamuri, William J. Gammerdinger, Reade B. Roberts, Karen L. Carleton, Sigbjørn Lien, Thomas D. Kocher

**Author notes:** Corresponding author orcid.org/0000-0002-7547-0133.

## Abstract

**Background:** African cichlid fishes are well known for their rapid radiations and are a model system for studying evolutionary processes. Here we compare multiple, high-quality, chromosome-scale genome assemblies to understand the genetic mechanisms underlying cichlid diversification and study how genome structure evolves in rapidly radiating lineages.

**Results:** We re-anchored our recent assembly of the Nile tilapia (*Oreochromis niloticus*) genome using a new high-density genetic map. We developed a new *de novo* genome assembly of the Lake Malawi cichlid, *Metriaclima zebra*, using high-coverage PacBio sequencing, and anchored contigs to linkage groups (LGs) using four different genetic maps. These new anchored assemblies allow the first chromosome-scale comparisons of African cichlid genomes.

Large intra-chromosomal structural differences (~2-28Mbp) among species are common, while inter-chromosomal differences are rare (< 10Mbp total). Placement of the centromeres within chromosome-scale assemblies identifies large structural differences that explain many of the karyotype differences among species. Structural differences are also associated with unique patterns of recombination on sex chromosomes. Structural differences on LG9, LG11 and LG20 are associated with reductions in recombination, indicative of inversions between the rock- and sand-dwelling clades of Lake Malawi cichlids. *M. zebra* has a larger number of recent transposable element (TE) insertions compared to *O. niloticus*, suggesting that several TE families have a higher rate of insertion in the haplochromine cichlid lineage.

**Conclusion:** This study identifies novel structural variation among East African cichlid genomes and provides a new set of genomic resources to support research on the mechanisms driving cichlid adaptation and speciation.

## Background

African cichlid fishes, due to their phenotypic diversity and rapid speciation over the last several million years, are a model system for studying the mechanisms of evolution [1]. Many studies of cichlid speciation have used short read data to perform genome scans of SNPs and small indels in order to identify genomic regions under selection [2–4]. However, there are numerous other ways that genomes can evolve, including the accumulation of larger indels, as well as intra- and inter-chromosomal rearrangements. Identification of these types of mutation requires high quality, nearly complete genome sequences.

Draft genomes of five African cichlid species were previously generated using Illumina short-read sequencing and used in an initial analysis exploring some of the forces at play in African cichlid speciation [5]. The draft genome assembly of the Lake Malawi cichlid, *Metriaclima zebra*, was one of the most continuous and accurate genomes assembled from short reads, as revealed in the Assemblathon 2 competition [6]. However, these five draft genome assemblies still contain a large number of gaps, and only the assembly of the Nile tilapia, *Oreochromis niloticus*, has been anchored to linkage groups (LGs), making it difficult to compare the structure of cichlid genomes at chromosomal scales.

To improve these cichlid genome resources, we have employed long-read Pacific Bioscience SMRT sequencing. Long-read DNA sequencing technology has made it much easier to create accurate and contiguous genome assemblies [7–11]. In particular, long-read technologies have allowed the assembly of repetitive sequences, and the identification of structural variants. We previously improved the genome assembly for the Lake Malawi cichlid, *M. zebra*, by sequencing 16.5X coverage of PacBio reads to fill in gaps and characterize repetitive sequences [12]. We also produced a new high-quality genome assembly of *O. niloticus*, using 44X coverage PacBio sequencing. We were able to anchor 86.9% of the assembly to linkage groups, which allowed us to characterize the structure of two sex determination regions in tilapias [13].

Cichlid karyotypes are similar to other perciform fish. The diploid chromosome number (2n) varies from 32-60, but more than 60% of species have a diploid number of 2n = 48 [14]. Most of the chromosomes are acrocentric, but between 0 and 9 metacentric pairs are present in each species [15,16]. These karyotypic changes may have played an important role in the evolution and speciation of African cichlids. Classical cytogenetic techniques are able to characterize differences in chromosome number and large fusion or translocation events, which are easily seen under the microscope. They are less suited to studying smaller genome rearrangements, including inversions smaller than several megabases. Comparisons of chromosome scale assemblies in other vertebrate groups have begun to identify extensive structural differences at both the cytogenetic and the sequence assembly level [17,18], but the role of chromosome rearrangements in recent adaptive radiations has not been well studied.

Chromosome-scale assemblies can be achieved either by physical mapping techniques [19], or by anchoring the contigs of the sequence assembly to genetic linkage maps. Genetic maps have the advantage of reflecting another important feature of genomes, namely variation in recombination rate, which has manifold impacts on the levels of genetic polymorphism [20] and the efficiency of genome scans [21].

Here we describe chromosome-scale assemblies of two cichlid genomes. First, we re-anchor our previously published PacBio assembly of the *O. niloticus* genome [13] using a new high-density genetic map [22]. Second, we present a new assembly of *M. zebra* based on 65X coverage of long PacBio sequence reads. Finally, we anchor the *M. zebra* assembly with several recombination maps produced from hybrid crosses among closely related species from Lake Malawi. The anchored genome assemblies of these two species allow for the first chromosome-scale comparison of African cichlid genomes. We focus our analyses on three aspects of genome evolution that are revealed by these new chromosome-scale assemblies: variation in recombination rate across the genome, structural variation among cichlid lineages, and the landscape of transposable elements.

First, we describe the pattern of recombination along each chromosome. Spatial variation in recombination rate has implications for patterns of genetic variation [23,24], the evolution of sex chromosomes [25], and the analysis of genome-wide associations between phenotypes and genotypes [21]. Despite the importance of recombination in shaping genome architecture [26], it is only beginning to be studied in cichlids [27]. A great diversity of sex chromosomes have evolved in East African cichlids, likely the result of sexual genetic conflict [28]. Rapid changes in sex determination mechanism, which are frequently variable even within species, may play an important role in cichlid speciation [1]. The evolution of new sex chromosomes often involves chromosomal inversions, which change the pattern of recombination [29–33]. Studies of these changing patterns of recombination, and their effects on genetic variation, have been hampered by the incomplete nature of the previous draft genome assemblies.

Second, we characterize the patterns of chromosome rearrangement among species. It has been suggested that teleost karyotypes have remained largely stable since the fish-specific whole genome duplication more than 300 million years ago [34]. This is in contrast to recent reports of chromosomal fusions among closely related cichlid species [35–37], and a large number of putative inversions associated with the evolution of sex chromosomes in various species [13,31,38]. Chromosome-scale assemblies of cichlids allow us to quantify the levels of synteny among teleost lineages, and the rate of intra-chromosomal rearrangement among cichlid lineages in East Africa. To further explore these distinct patterns of recombination and structural changes in cichlids, we also compared the cichlid genomes to the detailed genomic history of the medaka (*Oryzias latipes*). Previous studies in medaka have shown that, subsequent to the teleost-specific whole-genome duplication 320-350 million years ago, one subset of medaka chromosomes remained stable while another subset underwent more extensive fusion and translocation events [34,39]. Related comparisons using additional teleost species have shown that the diploid number of chromosomes is relatively stable (24-25 chromosomes pairs in 58% of teleosts) and that when the chromosome number is lower in a particular species or group it is due to chromosome fusion events [40].

Finally, we quantify the abundance and distribution of various transposable element (TEs) families in each genome. Several studies have documented the expansion of particular transposon families in East African cichlids (AFC TEs) [41,42]. Transposable elements may play an important role in shaping genome architecture, particularly the divergence of sex chromosomes. Transposable elements may also be an important source of regulatory mutations [43]. Insertion of an AFC-SINE into a gene promoter is associated with the evolution of a novel egg-spot coloration pattern in haplochromine cichlids [44]. Similar promoter element re-wiring events have been shown to control cichlid opsin visual sensitivity [45]. Since transposons may have been involved in the evolution of many other phenotypes, it is important that these sequences be well-represented in genome assemblies. Unfortunately, transposable elements are not well-represented in genome assemblies that are based on short Illumina sequence reads. Our previous work has shown that long-read sequencing greatly improves both the length and quantity of TE repeats in cichlid genome assemblies [12,13]. A comparative analysis of transposable elements will improve our understanding of the patterns of transposon insertion and deletion during the radiation of East African cichlids.

In this paper, we integrate the whole genome alignments of these two African cichlid species with the location of centromeres, and the patterns of recombination and linkage disequilibrium (LD). This panoramic view allows us to see how the structure of African cichlid genomes has evolved over the last several million years. It also raises new questions about the evolution of cichlid genomes, particularly about the evolution of sex chromosomes and the adaptive radiation of East African cichlids.

## Data Description

To begin this study of chromosome-scale comparisons of African cichlid genomes, we used a new high-density map of *O. niloticus* [22] to improve the anchoring of our recent genome assembly [13]. We also generated a high-quality *M. zebra* genome assembly from a single male caught on Mazinzi Reef in Lake Malawi. Single-molecule PacBio sequencing was performed to 65X coverage and a *de novo* assembly of the reads was constructed. Two new, and two previously published, genetic maps were used to quality check the assembly, break misassembled contigs, and anchor the sequence contigs to chromosomes. These new anchored genome assemblies of *O. niloticus* and *M. zebra* were then aligned to one another to compare their structure. The *O. niloticus* anchored assembly and sequencing reads are available under NCBI BioProject PRJNA344471. The *M. zebra* anchored assembly and sequencing reads are available under NCBI BioProject PRJNA60369.

## Analyses

### Anchoring the *O. niloticus* assembly to a high-density linkage map

The recently assembled *O. niloticus* genome [13] was re-anchored in this study using a new high-density map that includes 40,190 SNP markers, see Methods and [22]. This new map identified 22 additional misassemblies not identified by previous maps. Table 1 provides a comparison of the previous O_niloticus_UMD1 assembly with this newly anchored O_niloticus_UMD_NMBU assembly.

**Table 1.** Anchoring comparison of O_niloticus_UMD1 and O_niloticus_UMD_NMBU.

| Linkage group | O_niloticus_UMD1 | O_niloticus_UMD_NMBU | Change |
| --- | --- | --- | --- |
| LG1 | 38,372,991 | 40,673,430 | 2,300,439 |
| LG2 | 35,256,741 | 36,523,203 | 1,266,462 |
| LG3 | 68,550,753 | 87,567,345 | 19,016,592 |
| LG4 | 38,038,224 | 35,549,522 | -2,488,702 |
| LG5 | 34,628,617 | 39,714,817 | 5,086,200 |
| LG6 | 44,571,662 | 42,433,576 | -2,138,086 |
| LG7 | 62,059,223 | 64,772,279 | 2,713,056 |
| LG8 | 30,802,437 | 30,527,416 | -275,021 |
| LG9 | 27,519,051 | 35,850,837 | 8,331,786 |
| LG10 | 32,426,571 | 34,704,454 | 2,277,883 |
| LG11 | 36,466,354 | 39,275,952 | 2,809,598 |
| LG12 | 41,232,431 | 38,600,464 | -2,631,967 |
| LG13 | 32,337,344 | 34,734,273 | 2,396,929 |
| LG14 | 39,264,731 | 40,509,636 | 1,244,905 |
| LG15 | 36,154,882 | 39,688,505 | 3,533,623 |
| LG16 | 43,860,769 | 36,041,493 | -7,819,276 |
| LG17 | 40,919,683 | 38,839,487 | -2,080,196 |
| LG18 | 37,007,722 | 38,636,442 | 1,628,720 |
| LG19 | 31,245,232 | 30,963,196 | -282,036 |
| LG20 | 36,767,035 | 37,140,374 | 373,339 |
| LG22 | 37,011,614 | 39,199,643 | 2,188,029 |
| LG23 | 44,097,196 | 45,655,644 | 1,558,448 |
| Total anchored (%) | 868,591,263 (86.0%) | 907,601,988 (90.2%) | 39,010,725 (4.2%) |

The previous O_niloticus_UMD1 assembly anchored a total of 868.6Mbp while the new O_niloticus_UMD_NMBU assembly anchored a total of 907.6Mbp (90.2%). Much of the newly anchored sequence is on LG3, which increased by 19Mbp, from 68.6Mbp to 87.6Mbp. In the O_niloticus_UMD_1 assembly, LG3 was broken into LG3a and LG3b. The new O_niloticus_UMD_NMBU assembly merged these into a single LG3. LG3 is the largest and most repetitive chromosome in *O. niloticus* [15], and is a sex chromosome in the closely related species, *O. aureus* [46]. The anchored assembly of LG3 is 54.7% repetitive, compared to repeat rate of 37% genome-wide. The repetitive nature of *O. niloticus* LG3 is also highlighted by the fact that it required this new dense map to anchor many small contigs to this linkage group. Several chromosomes (e.g. LG16) have fewer total bp anchored in the O_niloticus_UMD_NMBU assembly compared to the previous O_niloticus_UMD1 assembly. This is due to the fact that misassembled contigs that have been broken by the new map are now assigned to a different LG.

### Diploid sequence assembly of *Metriaclima zebra*

We assembled 65X coverage PacBio reads using FALCON/FALCON-unzip [7] to generate the new diploid *M. zebra* assembly, “M_zebra_UMD2”. FALCON first assembles the PacBio reads into primary contigs (p-contigs) and unphased associate contigs (a-contigs) that correspond to alternate alleles. During the FALCON-unzip step, reads are assigned to haplotypes by phasing of heterozygous SNPs and then a final set of p-contigs and phased haplotigs are produced. Table 2 provides the assembly summary statistics for each of these assembly parts. The length of the p-contigs (total size 957Mb), compared to the estimated cichlid genome size of 1Gbp [47], suggests the assembly is relatively complete. To measure the completeness of the haplotigs, the theoretical sizes of heterozygous regions under null expectations of recombination rates and effective population sizes were compared to the size distribution of the haplotigs. Additional File A shows the size distribution of the assembled haplotigs and how it relates to the theoretical recombination rate for several different effective population sizes (N_e_). The shape of this haplotig size distribution is closest to the curves representing effective population sizes of 1,000-2,500, which closely matches a recent estimate of the effective population size in *M. zebra* [48]. Variance in recombination rate across the genome may bias this estimate.

**Table 2.** FALCON assembly results for *M. zebra*. NG50 and LG50 are based on an estimated genome size of 1Gbp [47]. N50 and L50 sizes are provided for a-contigs and haplotigs since the size for the alternate haplotype is not known.

| Assembly fraction | Assembly size (Mbp) | Number of contigs | NG50<br>N50 (Mbp) | LG50<br>L50 | Mean contig size (kbp) | Max contig size (Mbp) |
| --- | --- | --- | --- | --- | --- | --- |
| FALCON p-contigs | 986.67 | 3931 | 1.38 | 200 | 251.00 | 10.04 |
| FALCON a-contigs | 261.12 | 5625 | 0.054 | 1615 | 46.42 | 0.381 |
| FALCON-unzip p-contigs | 957.01 | 2313 | 1.42 | 186 | 413.75 | 10.01 |
| FALCON-unzip haplotigs | 642.33 | 6367 | 0.214 | 891 | 100.89 | 1.17 |

### Anchoring the *M. zebra* genome assembly

Four genetic recombination maps were used to detect misassemblies, anchor the contigs to chromosomes, and compare species level structural differences. The four maps were all produced from interspecific F2 crosses genotyped with RADseq strategies and involve six Lake Malawi cichlid species in total. The two previously generated maps were estimated using 160 F2 from a cross of *Metriaclima zebra* and *Metriaclima mbenjii* [49] and 262 F2 from a cross of *Labeotropheus fuelleborni* and *Tropheops* ‘*red cheek*’ [50]. The two new maps consisted of crosses of *M. mbenjii* x *A. koningsi* (331 F2) (*in preparation*) and *M. mbenjii* x *A. baenschi* (161 F2) (*submitted*). Table 3 provides the total bp anchored to each LG for each of the four maps. The final M_zebra_UMD2 assembly anchors 760.7Mbp.

**Table 3.** Anchoring of the *M. zebra* assembly to four different genetic linkage maps. The FALCON assembly was anchored to each map separately, and the total bases anchored are shown for each LG and map. The anchored map LGs that were used for the M_zebra_UMD2 anchoring are indicated in **<u>bold</u>**. The *L. fuelleborni x Tropheops* ‘*red cheek*’ map had four LGs that were combined into two (LG10a/LG10b and LG13a/LG13b). Selection of particular linkage groups for the final anchoring is based on accuracy and not necessarily overall length.

| Linkage group | *M. zebra* x<br>*M. mbenjii*<br>(160 F2) | *L. fuelleborni*<br>x *Tropheops*<br>'red cheek'<br>(262 F2) | *M. mbenjii* x<br>*A. koningsi*<br>(331 F2) | *M. mbenjii* x<br>*A. baenschii*<br>(161 F2) | M_zebra_UMD2 |
| --- | --- | --- | --- | --- | --- |
| LG1 | 31,191,433 | 32,150,205 | **38,662,702** | 36,192,366 | 38,662,702 |
| LG2 | 25,783,542 | 28,952,651 | **32,647,892** | 33,362,328 | 32,647,892 |
| LG3 | 18,498,838 | 14,707,016 | **37,717,145** | 24,847,713 | 37,309,556 |
| LG4 | 28,418,370 | 24,424,243 | **29,889,472** | 23,743,562 | 30,507,480 |
| LG5 | 29,725,229 | 34,008,850 | **36,154,892** | 30,984,548 | 36,154,892 |
| LG6 | 15,868,181 | 32,717,361 | **39,879,506** | 32,438,073 | 39,760,669 |
| LG7 | 29,333,014 | 57,016,972 | **64,381,187** | 50,973,986 | 64,889,811 |
| LG8 | 19,307,854 | 16,999,744 | **24,280,574** | 18,082,738 | 23,959,896 |
| LG9 | **21,018,370** | 22,620,859 | 18,771,712 | 24,011,483 | 21,018,370 |
| LG10 | 25,942,318 | 26,176,893 | **32,583,833** | 25,149,136 | 32,346,187 |
| LG11 | **32,253,887** | 30,903,800 | 34,404,464 | 31,577,152 | 32,434,411 |
| LG12 | 23,231,402 | 31,401,442 | **34,043,602** | 31,595,605 | 34,077,077 |
| LG13 | 25,893,161 | 24,034,634 | **31,886,878** | 28,831,406 | 32,061,881 |
| LG14 | 32,750,971 | 32,025,991 | **37,909,455** | 30,978,148 | 37,855,742 |
| LG15 | 28,015,059 | 28,462,857 | **34,537,245** | 28,405,563 | 34,537,245 |
| LG16 | 24,665,172 | 26,935,058 | **34,727,877** | 29,158,962 | 34,727,877 |
| LG17 | 28,473,329 | 31,631,813 | **35,766,785** | 31,607,415 | 35,766,785 |
| LG18 | 19,927,984 | 23,757,304 | **29,457,134** | 30,047,761 | 29,494,144 |
| LG19 | 24,076,222 | 19,992,035 | **25,739,093** | 22,726,673 | 25,955,740 |
| LG20 | 28,281,247 | 30,800,769 | 24,975,175 | **29,774,176** | 29,774,176 |
| LG22 | 27,460,019 | 31,372,369 | **34,717,234** | 30,512,954 | 34,717,234 |
| LG23 | 27,069,552 | 27,967,022 | **42,736,004** | 37,848,175 | 42,076,657 |
| Total anchored<br>(%) | 567,185,154<br>(59.3%) | 629,059,888<br>(65.7%) | 755,869,861<br>(79.0%) | 662,849,923<br>(69.3%) | 760,736,424<br>(79.5%) |
| Total including<br>unanchored | 957,158,042 | 957,163,242 | 957,185,442 | 957,167,042 | 957,200,631 |

Prior to the final anchoring, these four maps were also used to detect and confirm potential misassemblies in the FALCON contigs. Additional File B lists the FALCON p-contigs for which markers from two or more different LGs aligned, an indicator of potential inter-LG misassembly. Each of these potential misassemblies was further evaluated using alignments of a 40kb Illumina mate-pair library [5], RefSeq gene annotations [51], and repeat annotations (see Methods). In some cases, it was determined that the map marker sequences were repetitive, giving a false signal of misassembly. A total of 33 potential misassemblies were inspected and 16 likely misassemblies were identified and broken. An example of one of these misassemblies is provided in Additional File C. Whole genome alignment comparisons (see section below) detected one additional intra-chromosomal misassembly, bringing the final total to 17 misassemblies.

The *M. mbenjii* x *A. koningsi* map typically anchored more of the *M. zebra* assembly contigs, and in a more accurate order (i.e. greater collinearity with *O. niloticus*), than did the other three maps. This is likely due to the fact that the *M. mbenjii* x *A. koningsi* map had both more F2 individuals and more map markers than the other three Lake Malawi cichlid maps, giving it the highest resolution. Anchoring with the other three maps resulted in anchoring of more contigs on LG2, LG9, LG18, LG20 (see Table 3). However, the map that produced the longest anchored LG did not always appear to be the most accurate. To determine this accuracy, each *M. zebra* LG (anchored with each of the four maps) was aligned to the anchored *O. niloticus* assembly and compared (Additional File D). In the final assembly, the *M. zebra* x *M. mbenjii* map was used to anchor LG9 and LG11, and the *M. mbenjii* x *A. baenschi* map was used to anchor LG20. The anchoring of LG9 using *M. mbenjii* x *A. koningsi* map was very short compared to the other LGs and the other three maps anchored significantly more of LG9. The *M. zebra* x *M. mbenjii* map was chosen to anchor LG9 as it showed the most similar ordering relative to the *O. niloticus* assembly (Additional File D). The *M. zebra* x *M. mbenjii* map was also chosen to anchor LG11 as the other three maps showed large putative structural differences (Additional File D and also seen in the recombination maps, presented below). LG20 was best represented by the *M. mbenjii* x *A. baenschi* map based on alignment to *O. niloticus*, overall size and by ordering of markers in the recombination maps. Thus, the final M_zebra_UMD2 anchoring used three of the four maps to assign, order and orient contigs. The *L. fuelleborni x Tropheops ‘red cheek’* map was not used in the final anchoring but helped confirm many misassemblies and provided information on structural differences. Several LGs have slightly different overall sizes than when the assembly was anchored with just a single map (e.g. LG3 changed from 37,717,154bp to 37,309,556bp, Table 2). This is due to the fact that several small contigs are assigned to different LGs by the four different maps.

An anchoring analysis that sequentially chained together the anchored assemblies from all four Lake Malawi cichlid maps resulted in a slightly more anchored assembly (833Mbp total compared to 760Mbp for M_zebra_UMD2). However, the ordering of contigs in this combined anchored assembly was far less accurate (when aligned to *O. niloticus*) and so it was not used. There was only a single contig longer than 1Mbp (“000254F”) that was not anchored by at least one map.

### Structural differences among Lake Malawi cichlid genomes

The process of anchoring the M_zebra_UMD2 assembly to the four maps also allowed us to look for large structural differences among the six species used to generate the maps. Specifically, we looked for p-contigs that were assigned to different LGs in any of the four maps. Table 4 provides the list of the 9 contigs that were assigned to different LGs by at least two maps and which represent putative inter-chromosomal rearrangements.

**Table 4.** Putative inter-chromosomal differences as identified by map anchoring comparison. The number of markers aligned to each contig for each LG is indicated in (*N*). ‘NA’ indicates that a particular map had no markers aligned to that contig.

| contig name | contig size | *Mz.* x *Mb.*<br>map LG | *Lf.* x *Tr.*<br>map LG | *Mb.* x *Ak.*<br>map LG | *Mb.* x *Ab.*<br>map LG | Notes |
| --- | --- | --- | --- | --- | --- | --- |
| 000084F_pilon\|quiver | 2,383,905 | LG1 (1) | LG3 (3) | LG3 (6) | LG3 (3) |  |
| 000105F_pilon\|quiver<br>_1_1312536 | 1,312,536 | NA | LG10a (1) | LG2 (1) | LG2 (3) |  |
| 000201F_pilon\|quiver | 1,489,552 | LG3 (1) | *LG1* (3) | LG3 (3) | LG3 (1) |  |
| 000223F_pilon\|quiver | 1,452,516 | LG8 (4) | LG8 (8) | LG3 (2) | LG8 (4) | repetitive<br>markers on<br>LG3 |
| 000256F_pilon\|quiver | 1,241,607 | LG20 (1) | LG20 (1) | NA | LG9 (1) |  |
| 000414F_pilon\|quiver | 805,874 | LG5 (1) | LG5 (1) | NA | LG3 (1) |  |
| 000521F_pilon\|quiver | 566,343 | LG15 (2) | NA | LG17 (1) | NA | repetitive<br>marker on<br>LG17 |
| 000541F_pilon\|quiver | 515,490 | NA | LG2 (1) | LG3 (1) | NA |  |
| 000671F_pilon\|quiver | 374,096 | LG23 (1) | NA | LG23 (1) | LG22 (1) |  |

Seven of these nine contigs are anchored to a different linkage group in one of the maps by only a single marker. It is difficult to determine if there is a true inter-chromosomal difference with such little evidence. Even when all nine contig anchoring differences are considered, it amounts to only 10.1Mbp of total inter-chromosomal differences between the species used to generate the maps. This suggests that at most 1% of these Lake Malawi cichlid genomes have inter-chromosomal rearrangements relative to the reference anchoring. It is possible that there are some other significant inter-chromosomal differences that we did not detect in the unanchored portion of the genome. If they do exist, they are likely to be highly repetitive portions of these genomes that could not be assembled into the long contigs that can be accurately anchored.

### Localization of centromeric repeats

The location of centromeres is key to understanding structural rearrangements in the karyotype. Figure 1 shows the karyotype of *O. niloticus* and *Metriaclima lombardoi* (a species closely related to *M. zebra*). The *O. niloticus* SATA consensus repeat [52] is common to the centromeres of many East African cichlid [15], and closely matches the satellite repeats identified in a recent analysis of centromeres across many taxa [53].

**Figure 1.**
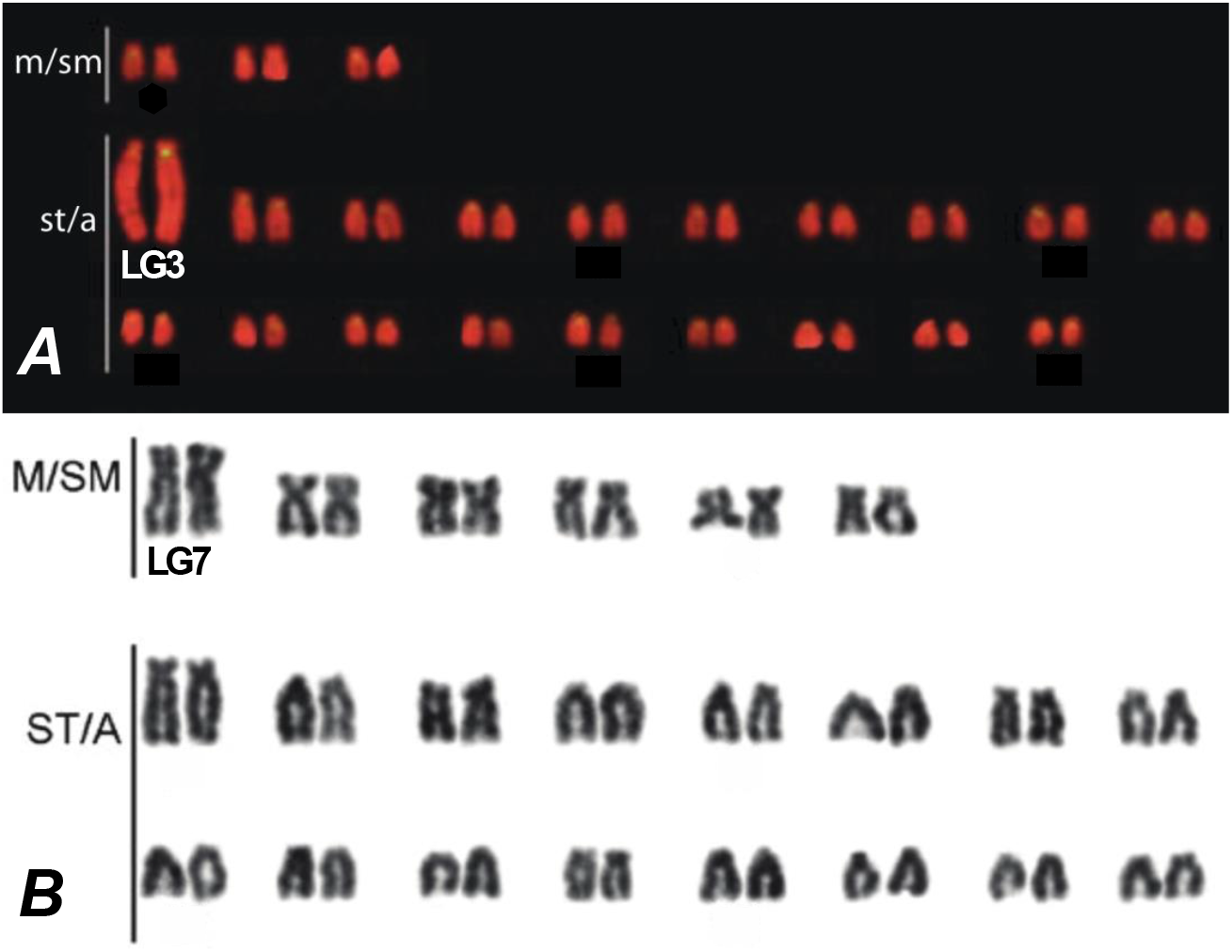
A) Chromosome mapping of SATA satellite DNA in *O. niloticus* reproduced and modified with permission from [15]. The SATA sequences are labelled in yellow against the background staining with propidium iodide. B) Giemsa-stained karyograms of the Lake Malawi *Metriaclima lombardoi* reproduced and modified with permission from [33]. LG3 in *O. niloticus* (A) and LG7 in *Metriaclima* (B) are labeled based on [36].

*Oreochromis and Metriaclima* diverged 17-28 million years ago [54]. Their karyotypes each have 22 chromosome pairs, as do the majority of African cichlids, but *O. niloticus* has 1 to 3 meta-submetacentric and 19 to 21 subtelo-acrocentric chromosomes according to two previous karyotypes [15,55], whereas *M. zebra* has six meta-submetacentric and 16 subtelo-acrocentric chromosomes. The chromosomes in Figure 1 have been ordered by type and then by size but only a few have been previously assigned to LGs. BAC and additional marker sequences have been used for specific labeling of chromosomes [36,56].

In order to understand the structural basis for these differences in karyotype, we constructed and visualized whole genome alignments of M_zebra_UMD2 and O_niloticus_UMD_NMBU (Additional File D). Figure 2 shows the LG23 alignment of *M. zebra* and *O. niloticus*. Placement of centromere repeats identify a large structural rearrangement on LG23 that shows that this chromosome is subtelo-acrocentric in *O. niloticus*, but meta-submetacentric in *M. zebra*.

**Figure 2.**
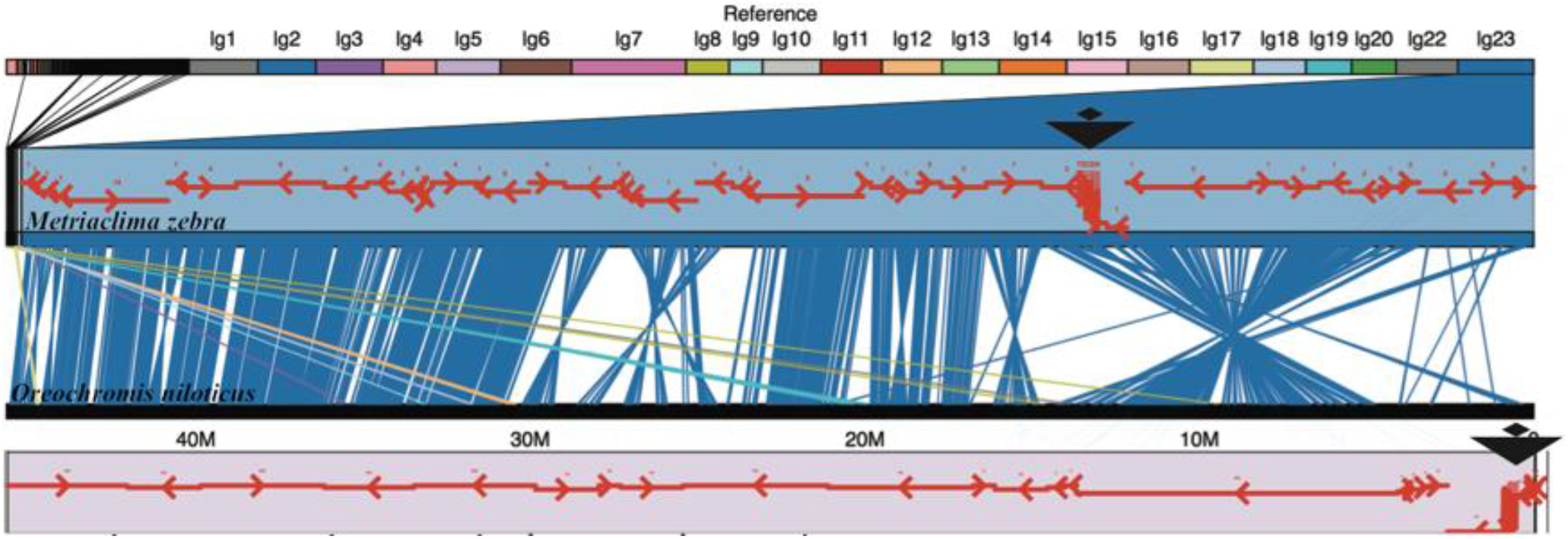
Comparative alignment of LG23 in *M. zebra* and *O. niloticus*. Centromere repeats in each assembly are indicated by large black triangles. Anchored contigs in each assembly are shown as red arrows indicating the orientation of each contig.

Centromere repeats were not assembled on every chromosome for either *M. zebra* and *O. niloticus*. However, on chromosomes where centromere repeats were placed in both assemblies, and a large structural difference was observed, we were able to identify large centromere repositioning events, including acrocentric/metacentric changes on LG4, LG7, LG16, LG17, and LG23. Although we were not able to identify the centromeres in both genome assemblies, similar rearrangement events suggest acrocentric/metacentric changes on LG2, LG6, LG20, and LG22 as well (Additional File D).

The whole genome alignment comparisons of *M. zebra* and *O. niloticus* also identified a number of large intra-chromosomal structural rearrangements that do not involve the centromere. On LG2 there are two large rearrangements of ~15Mbp and ~20Mbp (Additional File D). The largest single structural change appears on LG19 where there is a ~23Mbp rearrangement between *M. zebra* and *O. niloticus*. A similar ~20Mbp rearrangement is present on LG20. There is an ~11Mbp rearrangement at one end of LG22 that may be associated with another change in centromere location, although the centromere was not localized on LG22 in either assembly.

Perhaps the most diverged chromosome in terms of size, structure and repeat content is LG3. The karyotype of *O. niloticus* LG3 is much larger and more repetitive than the corresponding LG3 in Lake Malawi cichlids (Figure 1 and [15,55]). In the related species *O. aureus*, sex determination is controlled by a locus on LG3 [13,46]. Additional File E shows an F_ST_ comparison of the *O. aureus* male versus female pools described in [13]. There is a very wide region of sex-patterned differentiation in *O. aureus* on LG3 from ~40Mbp to 85Mbp. The large karyotype of LG3 in *O. niloticus* reflects both this large region of differentiation associated with the sex-determination locus as well as the vast amounts of repetitive sequence that have accumulated in this region.

### Variation in recombination rate among species

To compare the rates and patterns of recombination across the chromosomes, each set of map markers was aligned to the corresponding assembly and their recombination map positions plotted against physical distance. Male and female recombination in *O. niloticus* is plotted against the O_niloticus_UMD_NMBU assembly in Additional File F. Overall, the *O. niloticus* chromosomes are characterized by low recombination on the ends of chromosomes and higher recombination in the middle of chromosomes. Each of the *O. niloticus* chromosomes show a difference in recombination between males and females. The typical pattern is higher recombination in the females than the males. However, LG6 and parts of LG4, LG9, LG16, LG20, and LG22 show higher recombination in males than females. LG3 and LG23 are both known sex determination chromosomes in tilapias [46,57], and each deviates from the normal recombination patterns. On LG3, the largest chromosome in *O. niloticus* (Figure 1), there is very low recombination for ~70Mbp. On LG23 there is a ~28Mbp region of greatly reduced recombination.

Likewise, the markers in the four Lake Malawi genetic recombination maps were aligned to the final M_zebra_UMD2 assembly and their recombination map positions were plotted against physical distance. Figure 3 shows the comparison of the four Lake Malawi genetic recombination maps relative to the M_zebra_UMD2 anchored assembly for four chromosomes. Additional File G contains plots for the other chromosomes. Similar to the *O. niloticus* chromosomes, many Lake Malawi chromosomes show low recombination on the ends of chromosomes and higher recombination in the middle of chromosomes, with several notable exceptions that are indicative of structural changes. In the Lake Malawi maps (Additional File G) there is a region of low recombination for the first ~15Mb of LG2 that corresponds with a large structural rearrangement relative to *O. niloticus* (Additional File D). On LG7 (Figure 3) the usual pattern of low recombination at the ends of the chromosomes is observed in all four maps, but there is also a region of low recombination in the middle of the chromosome (at ~30Mbp in M_zebra_UMD2), near several smaller scale rearrangements relative to *O. niloticus* (Additional File D). An XY sex determination locus has been identified in this region of LG7 in many Lake Malawi species [29,58]. There is also evidence of large structural rearrangements on LG9 in all four Lake Malawi crosses, as evidenced by both the recombination map and whole genome alignment comparisons (Additional Files D and G). There appears to be a ~2Mbp inversion on LG10 (relative to *O. niloticus*) that is associated with lowered recombination around 20Mbp in M_zebra_UMD2 (Additional Files D and G). LG11 (Figure 3) follows the typical recombination pattern for the *M. zebra* x *M. mbenjii* map, but there appears to be a large 15Mbp inversion in the genus *Aulonocara*, inferred from a large region of complete recombination suppression found in both the *M. mbenjii* x *A. koningsi* and *M. mbenjii* x *A. baenschi* maps. This likely corresponds to another sex locus. The *L. fuelleborni* x *Tropheops* ‘*red cheek*’ map also shows a large, but different, rearrangement on LG11 when compared to *O. niloticus*. LG15 has a region of lower recombination in the middle that is also associated with structural rearrangements relative to *O. niloticus* (Additional Files D and G). There is a large structural rearrangement on LG20 present in each of the four anchored assemblies that is also associated with a large (~15Mbp) region of low recombination (Figure 3 and Additional File D). Each of the four maps shows high recombination from 0-15Mbp and then much lower recombination to the end of LG23, although the *M. zebra* x *M. mbenjii* map does not show as much reduction in recombination than the other three maps (Figure 3). The centromere of LG23 is placed at 30.1Mbp and is in the middle of the region of low recombination. During the process of anchoring the M_zebra_UMD2 assembly with these maps, one additional misassembly was detected at 6,922,000 on contig 000000F on LG12 and subsequently broken for the final anchoring (included in Additional File G).

**Figure 3.**
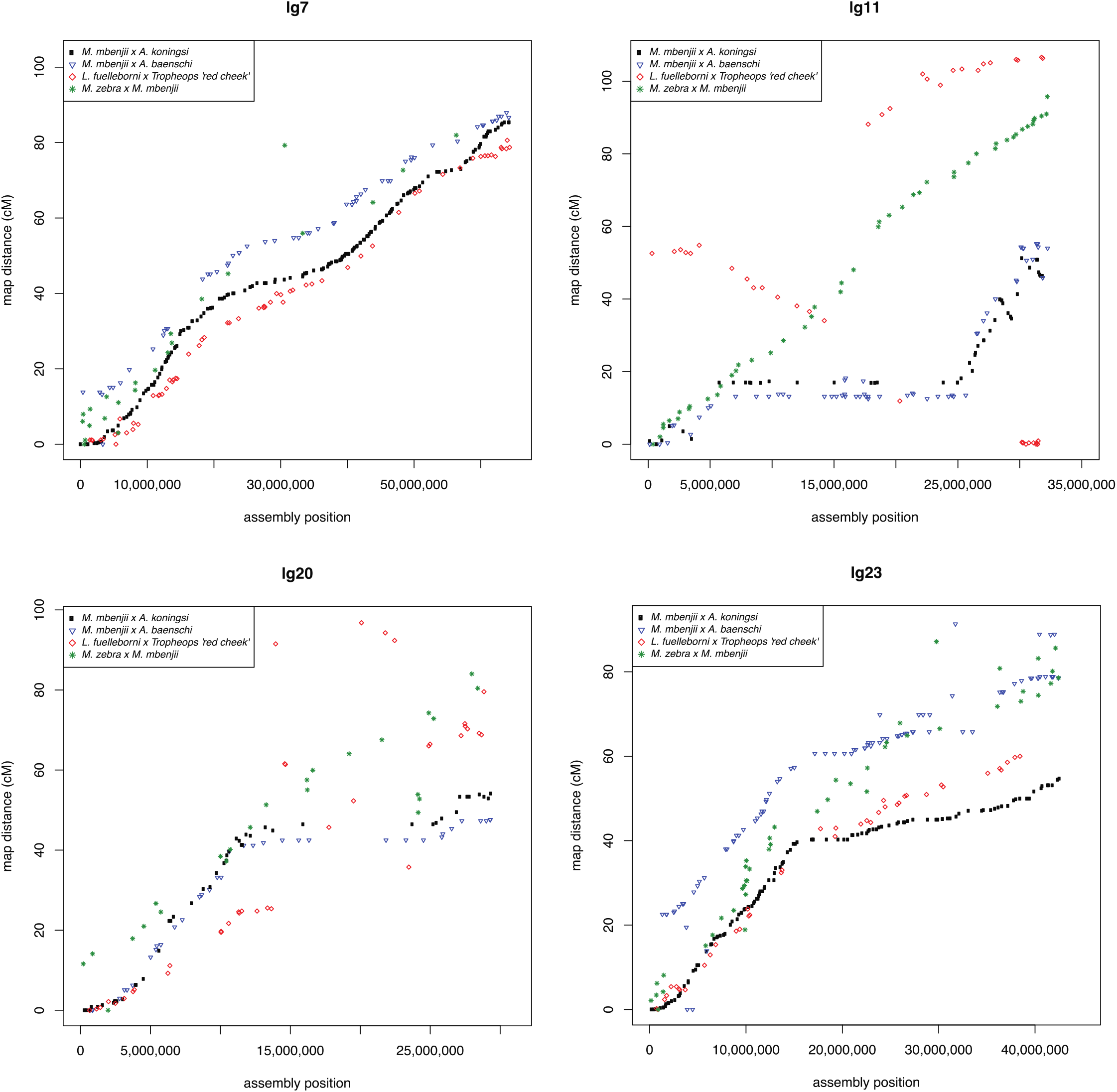
Comparison of the four maps relative to M_zebra_UMD2 on for LG7, LG11, LG20 and LG23. All LG maps are provided in Additional File G.

### Major structural rearrangements of ancient cichlid chromosomes

We also aligned the O_niloticus_UMD_NMBU assembly to the recently published “HSOK” *O. latipes* medaka assembly [39]. *O. niloticus* has 22 chromosome pairs, while the medaka HSOK genome has 24 chromosome pairs. Table 5 is a comparison of cichlid chromosomes and medaka HSOK chromosomes.

**Table 5.** Correspondence between *O. niloticus* and *O. latipes* chromosomes. Alignment lengths are provided for chromosomes with large fusion/translocation events.

| O_niloticus_UMD_NMBU chromosome | Primary medaka HSOK chromosome (alignment length) | Secondary medaka HSOK chromosome (alignment length) |
| --- | --- | --- |
| LG1 | 3 |  |
| LG2 | 10 |  |
| LG3 | 18 |  |
| LG4 | 8 |  |
| LG5 | 5 |  |
| LG6 | 1 |  |
| LG7 | 6 (32Mbp) | 12 (31Mbp) |
| LG8 | 19 |  |
| LG9 | 20 |  |
| LG10 | 14 |  |
| LG11 | 16 |  |
| LG12 | 9 |  |
| LG13 | 15 |  |
| LG14 | 13 |  |
| LG15 | 24 (31Mbp) | 4 (5Mbp) |
| LG16 | 21 |  |
| LG17 | 23 (23Mbp) | 4 (12Mbp) |
| LG18 | 17 |  |
| LG19 | 22 |  |
| LG20 | 7 |  |
| LG22 | 11 |  |
| LG23 | 2 (23Mbp) | 4 (17Mbp) |

We identified several large chromosome rearrangements that occurred in a cichlid ancestor. Tilapia LG7, the second largest chromosome (Table 1), is comprised of medaka chromosomes 6 and 12 in their entirety (Figure 4). This indicates a fusion of these ancestral chromosomes in cichlids relative to medaka, as had been previously suggested [37]. Tilapia LG23, the third largest chromosome (Table 1), is comprised of medaka chromosome 2 in its entirety and 17Mbp, or roughly half, of medaka chromosome 4 (Figure 5). The other half of medaka chromosome 4 was likely translocated onto LG15 and LG17. While the remaining 18 chromosomes have undergone extensive intra-chromosomal rearrangements, they have largely maintained a correspondence to individual medaka chromosomes over the course of the 120 million years of evolution since the last common ancestor of these species.

**Figure 4.**
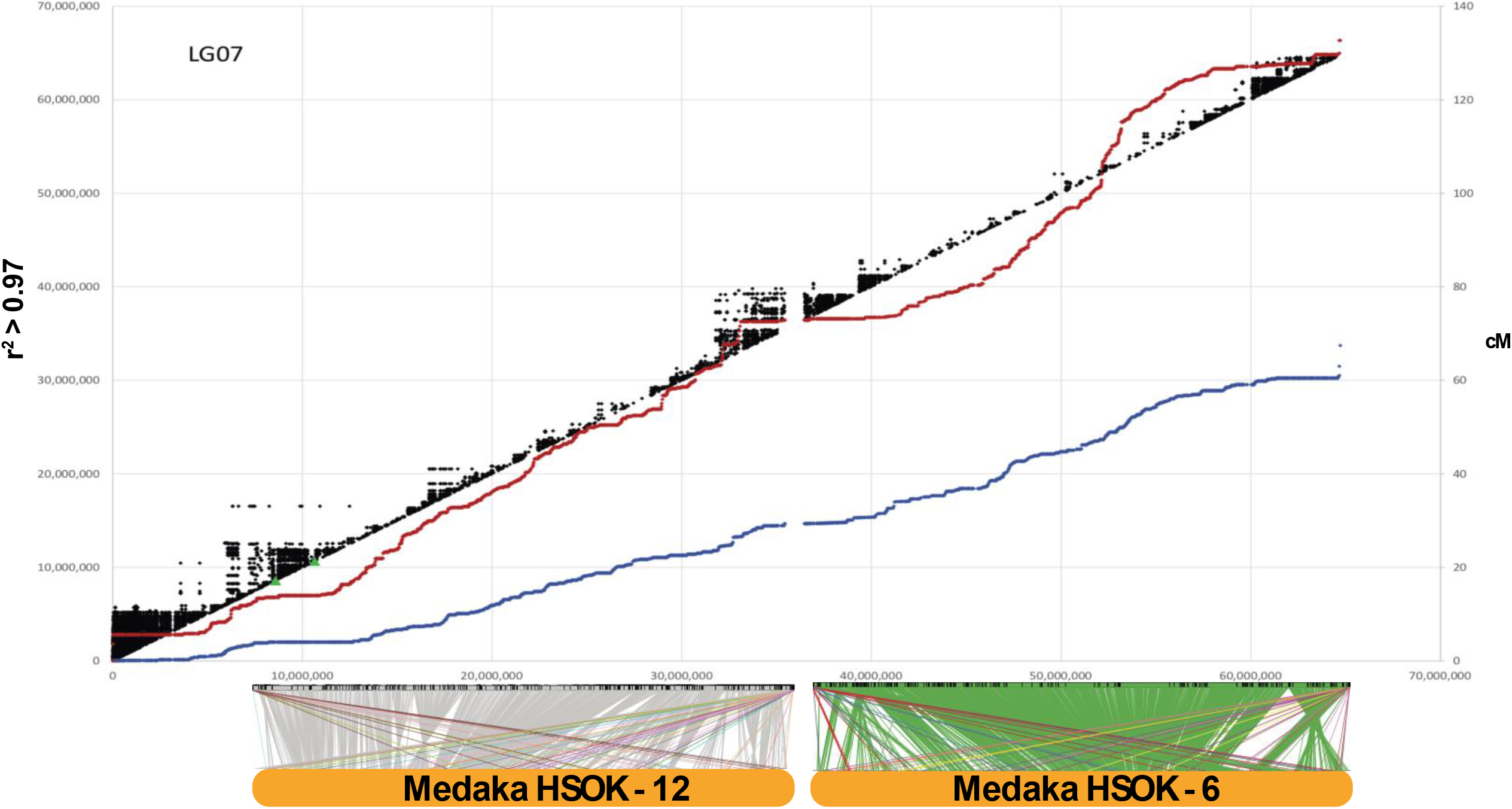
O_niloticus_UMD_NMBU LG7 is an ancient fusion of medaka HSOK 12 and 6. Female (red) and male *O. niloticus* recombination curves are shown along with linkage disequilibrium (r^2^ > 0.97) in black. Alignment of LG7 to medaka HSOK 12 and 6 are shown on the bottom.

**Figure 5.**
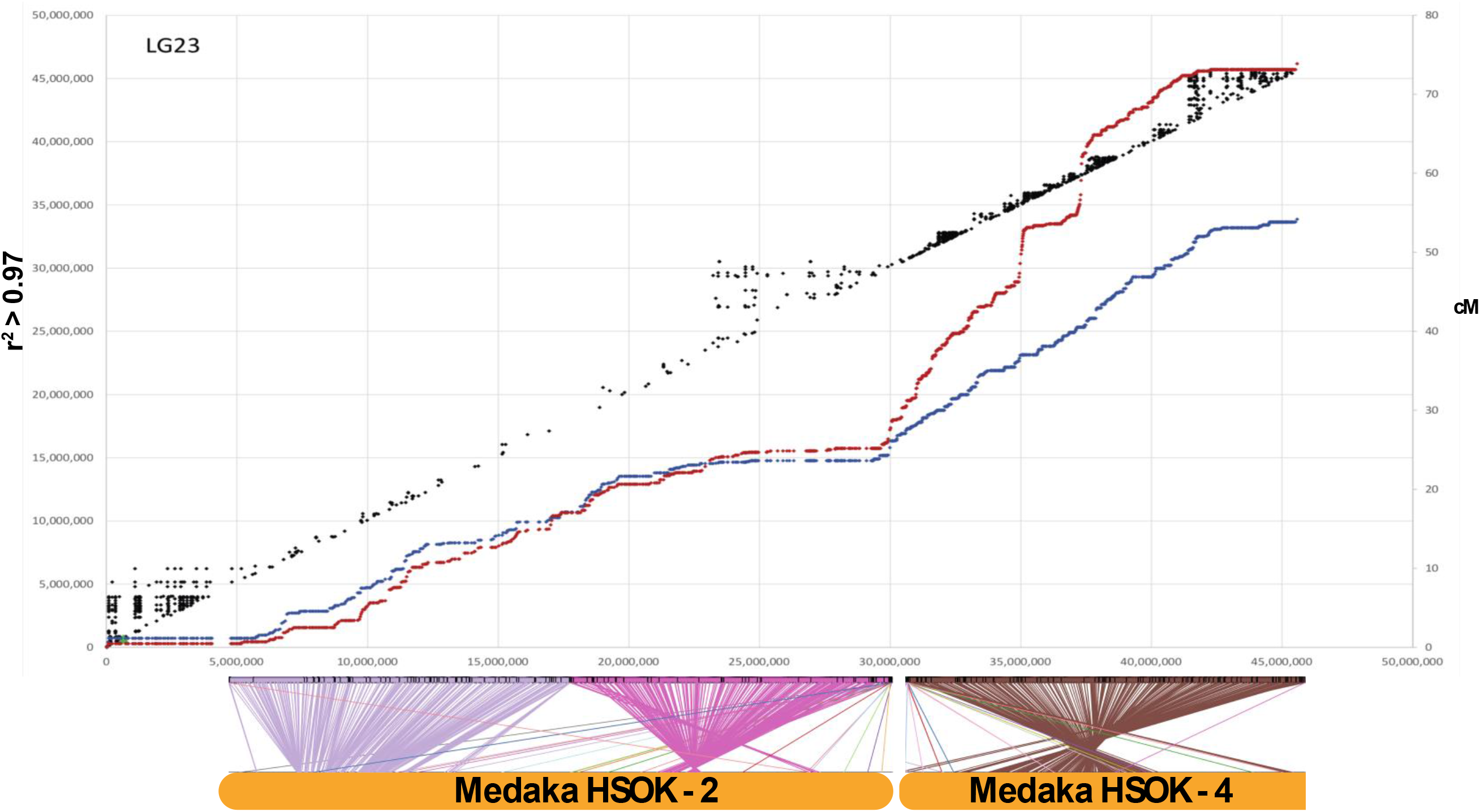
O_niloticus_UMD_NMBU LG23 is an ancient fusion of medaka HSOK 2 and part of medaka HSOK 4. Female (red) and male *O. niloticus* recombination curves are shown along with linkage disequilibrium (r^2^ > 0.97) in black. Alignment of LG7 to medaka HSOK 12 and 6 are shown on the bottom.

LG3 is the largest tilapia chromosome (Table 1), but surprisingly does not show any evidence of a chromosomal fusion or translocation event. Tilapia LG3 aligns well to medaka chromosome 18 along the first ~30Mbp of LG3, and the remainder of LG3 aligns to medaka chromosome 18 with much less contiguity. This divergent region of LG3 corresponds to the large, ~70Mbp region of low recombination in tilapia.

### Linkage disequilibrium

There is significant linkage disequilibrium (LD) in the tilapia GST^®^ population (see Methods), as shown in Figure 4 and Figure 5. As expected, the regions of low recombination near the ends of the chromosome show the highest levels of linkage disequilibrium. Large blocks of LD are also evident around the centromere on LG15 (Additional File F), and in the low recombination regions associated with the ancestral chromosome fusions on LG7 (Figure 4) and LG23 (Figure 5).

### Repeat landscape of the *Metriaclima zebra* assembly

The M_zebra_UMD2 assembly is 35% repetitive, similar to the O_niloticus_UMD1 assembly which is 37% repetitive [13]. Figure 6 shows the repeat landscape for the *M. zebra* and *O. niloticus* assemblies. While the *O. niloticus* genome assembly does have a slightly larger total quantity of repeats, the *M. zebra* genome assembly has a noticeably larger amount of recent TE insertions (sequence divergence < 2%). To further test that this difference was not an artifact of the two assembly processes, we assembled the *M. zebra* PacBio reads at the same coverage, with the same parameters, using the same software versions and on the same compute cluster as was performed for the O_niloticus_UMD1 assembly. RepeatMasker was subsequently run on this assembly and the pattern of more recent insertion in *M. zebra* relative to *O. niloticus* was even more pronounced (Additional File H).

**Figure 6.**
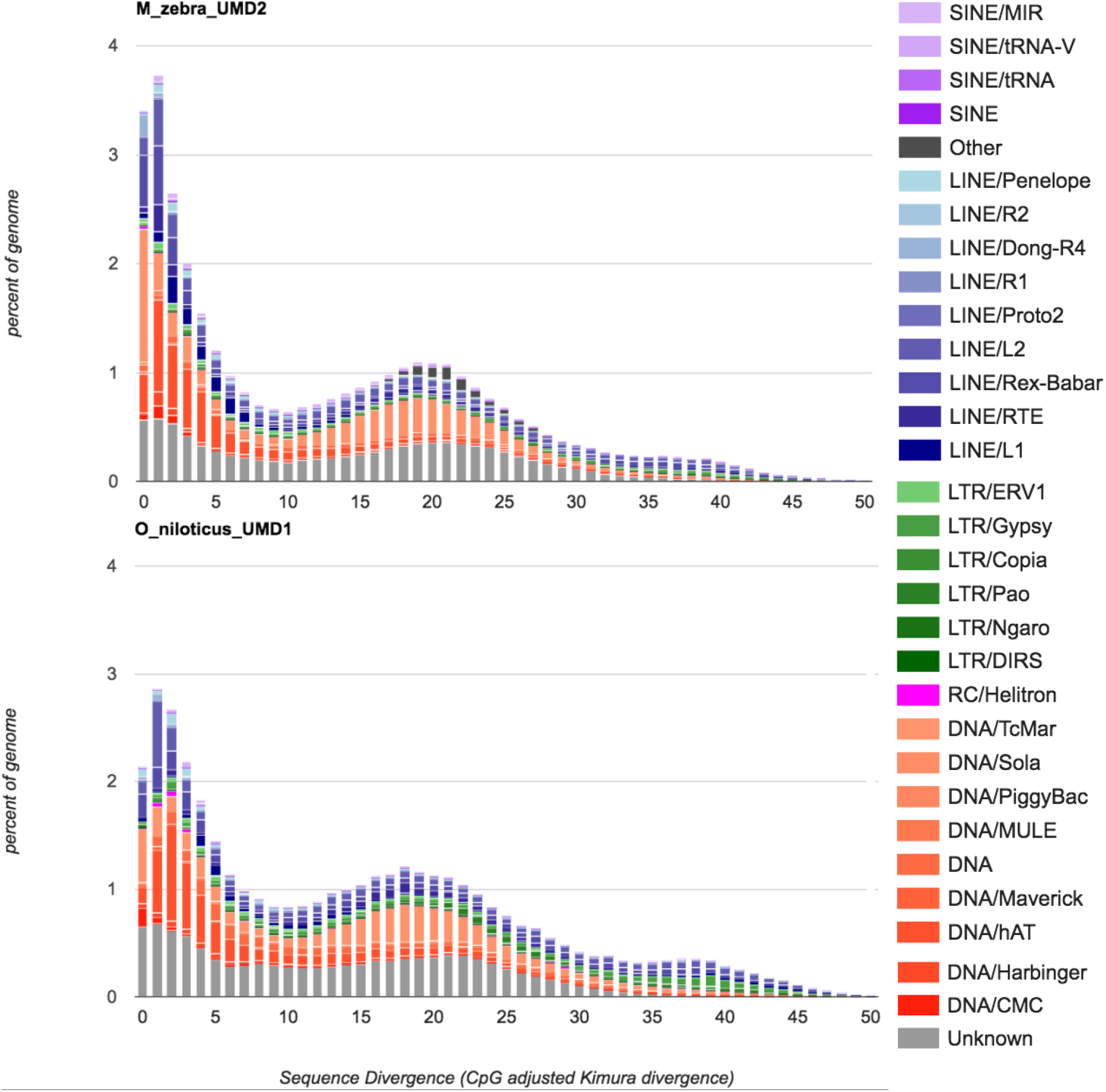
Comparison of the repeat landscape in the *M. zebra* and *O. niloticus* genome assemblies.

Three TE families account for most of the difference in the recent TE activity between the two species. Recent insertions (defined as 0-1% sequence divergence) of the class II DNA transposon superfamily, Tc1-Mariner, make up 0.5% of the total O_niloticus_UMD1 assembly, but make up 1.3% of the *M. zebra* assembly. Another class II DNA transposon superfamily, hAT, has recently inserted and makes up 0.15% of the O_niloticus_UMD1, but makes up 0.45% of the *M. zebra* assembly. Recent insertions of the class I retrotransposon superfamily, LINE-Rex-Babar, make up 0.2% of the O_niloticus_UMD1 assembly, but make up 0.6% of the *M. zebra* assembly. Other TE superfamilies show smaller increases in *M. zebra* as well. This indicates that *M. zebra*, and perhaps Lake Malawi cichlids in general, have experienced more recent TE expansion than the riverine counterpart, *O. niloticus*.

Overall, the amount of TEs assembled has increased from the original Illumina-only based *M. zebra* assembly [5], to the moderate PacBio coverage gap-filled M_zebra_UMD1 assembly [12], to the high PacBio coverage M_zebra_UMD2 assembly. Additional File I provides a comparison of repeat landscapes for each of these three *M. zebra* assemblies. The overall number of TEs, and particularly the most recently inserted TEs, are better represented as the assemblies improve. The African Cichlid-specific AFC-SINEs and AFC-LINEs [59], have been assembled in greater length as well. For example, the ~7.1kb “L1-1_AFC” LINE was assembled into 2,874 copies (across 1.29Mbp) in the original M_zebra_v0 assembly, 1,350 copies (across 1.66Mbp) in the M_zebra_UMD1 assembly and 2,295 copies (across 4.77Mbp) in the new M_zebra_UMD2 assembly.

### Genome completeness and annotation

Benchmarking Universal Single-Copy Orthologs (BUSCO) [60,61] was used to assess the completeness of the new *M. zebra* genome assembly. 2,586 complete vertebrate BUSCOs were searched and 2,465 (95.3%) complete BUSCOs were found, of which 71 (2.7%) were duplicated and 2,394 were single-copy. Only 82 (3.2%) were reported as fragmented, and just 39 (1.5%) BUSCOs were reported as missing.

The M_zebra_UMD2 assembly was annotated using the NCBI RefSeq annotation pipeline for eukaryotic genomes [51]. Table 6 shows the improvement in gene annotation for the new M_zebra_UMD2 assembly relative to the previous version of the *M. zebra* assembly [5,12].

**Table 6.**
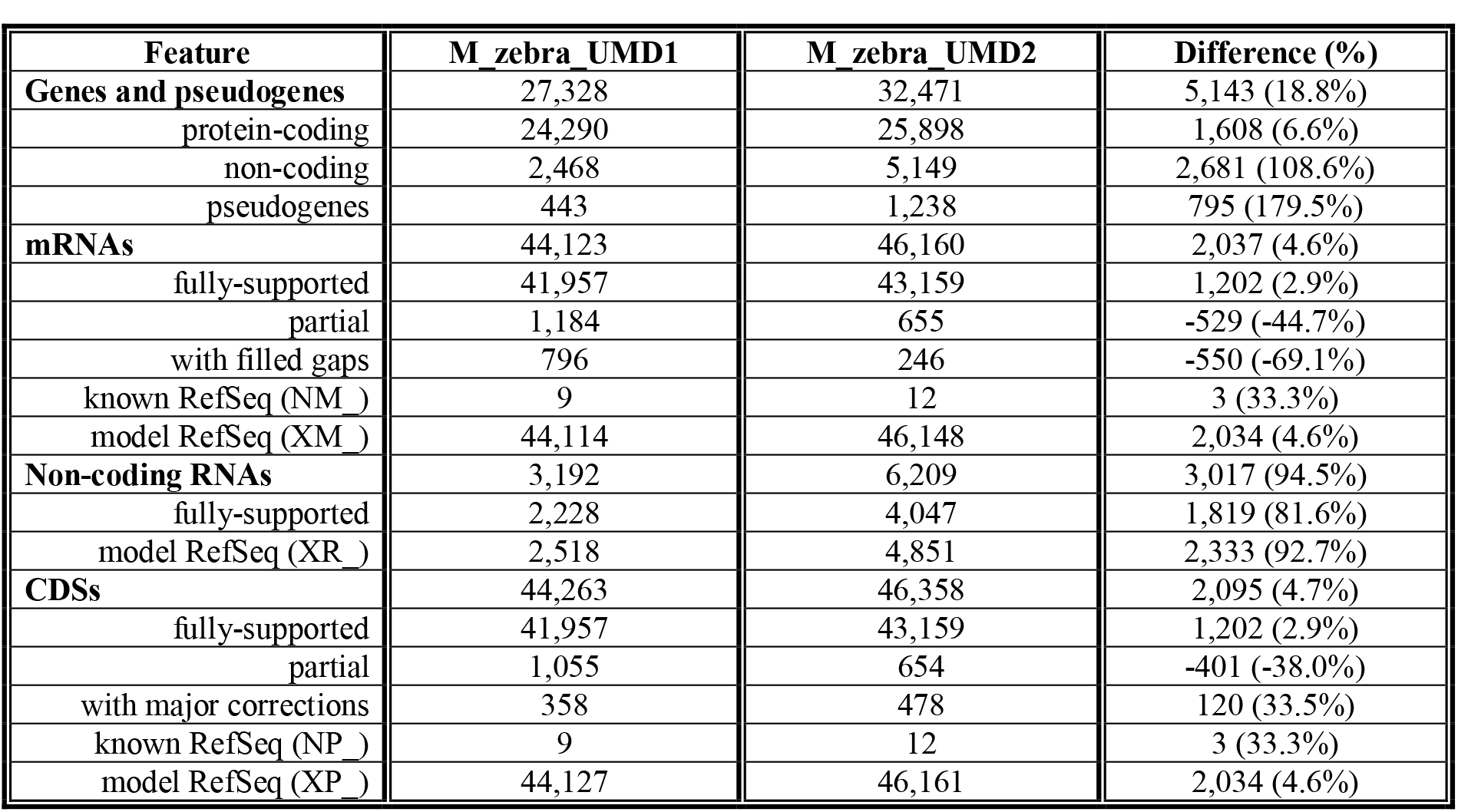
Annotation improvement of the M_zebra_UMD2 assembly gathered from RefSeq annotation reports [62,63].

## Discussion

### Anchoring to produce chromosome-scale assemblies

The genetic maps and whole genome alignment comparisons to the *O. niloticus* assembly were very useful for identifying large and mostly inter-chromosomal misassemblies in the new *M. zebra* assembly. A 40kb Illumina jumping library was also used in this process to determine if disagreements between the maps and the assembly were true misassemblies, errors in the maps, or structural differences between samples. It is likely that several misassemblies still remain in the final M_zebra_UMD2 anchoring. However, these potential misassemblies are probably only present on smaller contigs where there were not enough markers to detect misassembly events. Only one contig longer than 1Mbp was not able to be anchored by two or more markers from one of the four Lake Malawi maps. Therefore, any possible remaining misassemblies are likely to involve smaller contigs. The M_zebra_UMD2 anchoring was primarily anchored with the *M. mbenjii x A. koningsi* map, but for several chromosomes (Table 3), use of other Lake Malawi maps were used to arrive at a more complete anchoring. Therefore, the M_zebra_UMD2 anchoring does not represent a single species. A high-density map of *M. zebra* would be a useful resource for future studies.

### Patterns of continuity in genome assemblies

The longest contigs tend to be anchored in the middle of chromosomes and in regions where there is greater recombination. The ends of chromosomes, typically in regions of lower recombination, tend to have smaller contigs. Perhaps the clearest example of this is on LG13 (Additional File D and Additional File G). On LG7, smaller contigs appear in the middle of the chromosome where there is also a reduction in recombination uncharacteristic of most other chromosomes. Smaller contigs likely correspond to regions with a large fraction of repetitive sequence that lead to a more fragmented assembly. These regions have likely accumulated large TE arrays that are not spanned by even the longest of the reads in our datasets. It is known that TEs accumulate in regions of suppressed recombination [64]. These chromosomal regions with smaller contigs also tend to have more structural rearrangements relative to *O. niloticus*, which suggests an important role for transposable elements in formation of the rearrangements. This pattern could also be caused by ambiguities in the maps due to there being fewer recombination events and therefore less map resolution in these regions. There are also fewer markers used to anchor smaller contigs that may also contribute to this pattern. Orthogonal mapping technologies that do not rely on recombination, such as optical mapping, may be needed to resolve the structure of these regions in finer detail.

### Patterns of recombination in *O. niloticus*

Several patterns are evident in the recombination maps for *O. niloticus*. First, though the pattern of recombination is generally similar in males and females, the level of recombination in females is generally higher than in males. The total female map length is 1,641 cM, while the male map is only 1,321 cM. The sex differences in recombination rate of *O. niloticus* are smaller than observed in salmonids [65–68], stickleback [69], Japanese flounder [70], and zebrafish [71]. Second, the pattern of recombination on each chromosome is usually sigmoidal, with relatively little recombination over about 5Mb at the ends of each chromosome. The highest levels of recombination are found in the middle of each chromosome. This pattern is exactly opposite the pattern observed in stickleback and catfish, where recombination is highest at the ends of the chromosomes [72,73].

These patterns of recombination have implications for the pattern of linkage disequilibrium (LD) along each chromosome, which varies significantly across the genome. Blocks of LD are much longer in the regions of low recombination (Figure 4, Figure 5, Additional File F), such as near the ends of each chromosome. Regions of low recombination tend to accumulate repetitive transposable elements [64]. These regions are also likely to experience episodes of genetic hitchhiking, which will alter the pattern of genetic differentiation across the genome, as shown in stickleback [69,72]. The extent of LD impacts the probability of fixation of adaptive variants and may affect the probability that a given chromosomal segment can evolve into a new sex chromosome [69]. Interestingly, extensive LD is present on LG3 in *O. niloticus*, where a sex determination locus is located in a sister species, *O. aureus*. One evolutionary interpretation of this finding is that high LD at LG3 predated, and facilitated evolution of, the *O. aureus* sex chromosome [46]. Alternatively, recombination suppression may have evolved as a result of sex-chromosome-associated evolution at LG3; in this scenario, the lineage leading to *O. niloticus* may have had, and subsequently lost, the dominant LG3 sex determination allele, but the traces of sex chromosome evolution remains in the genome.

### Patterns of recombination in Lake Malawi cichlids

The four genetic maps of Lake Malawi cichlids show the same general pattern of recombination as *O. niloticus*. Again, the pattern of recombination on most Lake Malawi chromosomes is characterized by low recombination at the ends of the chromosomes and high recombination in the middle of the chromosomes. The several exceptions all indicate lineage-specific, intra-chromosomal rearrangements among the Lake Malawi species.

Perhaps the most striking difference between these four maps is a large (~19Mbp) putative inversion on LG11 in *Aulonocara*, as evidenced by the lack of recombination in the *M. mbenjii* x *A. koningsi* and *M. mbenjii* x *A. baenschi* maps (Figure 3). The *M. zebra* x *M. mbenjii* map does not contain this putative inversion and shows the normal recombination pattern across LG11, suggesting the inversion may represent an evolved difference between the *Metriaclima* and *Aulonocara* lineages. The *L. fuelleborni x Tropheops* ‘*red cheek*’ cross appears to show a different structural arrangement on LG11 but shows no evidence of suppressed recombination. There also appears to be a large inversion on LG20 in *Aulonocara* relative to the other three genera (Figure 3). Both of the *Aulonocara* maps show highly reduced recombination across a 15Mb region of this chromosome. All four crosses also showed a reduction in recombination for the first ~15Mbp on LG2 (Additional File G) that corresponds exactly with a structural rearrangement relative to *O. niloticus* (Additional File D). There is no such reduction in recombination on LG2 in *O. niloticus* (Additional File F), suggesting that the rearrangement occurred in the Lake Malawi ancestor and has maintained reduced recombination in this region across all lineages.

### Patterns of recombination on sex chromosomes

Sex chromosomes typically accumulate inversions that reduce recombination between the sex determining gene and linked sexually antagonistic alleles [74]. The strain of *O. niloticus* used to generate the genome assembly contigs [13] has an XY sex determination locus on LG1 [31,75]. The strain of *O. niloticus* used to generate the map [22] and anchor those contigs to chromosomes has an XY sex determination locus on [76]. We observed reduced recombination in males relative to females adjacent to the sex locus at 34.5Mbp on *O. niloticus* LG23 (Additional File F). We also observed significant differences in recombination between the sexes on LG7, LG11, LG14 and LG15. An XY sex locus has been identified on LG14 in *O. mossambicus* (Gammerdinger, *in review*), and XY sex loci have been identified on LG7 [29,58] and LG11 (unpublished) in Lake Malawi cichlids. Notably, three of the four Lake Malawi crosses segregate the LG7 XY sex determination system (the *M. mbenjii* x *A. baenschi* cross is unknown) (50,76 and unpublishsed results). However, LG7 shows relatively low recombination suppression compared to some other chromosomes. Recombination is reduced in the middle of LG7, centered at ~32Mbp, but this is not associated with the centromere (located at 61Mbp). While this region is near the LG7 XY sex determination interval, the overall shape of recombination on LG7 may also be the result of the chromosome fusion event that happened in the cichlid ancestor (Figure 4 and discussed below). As discussed for *Oreochromis* above, it is unclear whether recombination suppression or sex determination evolved first at this locus. It should also be noted that there is a single marker in this region that appears out of order in the *M. zebra* x *M. mbenjii* map, perhaps indicating a structural difference (Additional File D and Figure 3). Further investigation will be needed to determine if other regions of the genome that display large differences in sex-specific recombination are associated with previously identified and/or novel sex determination loci.

The new anchoring provides the most complete assembly to date of LG3, the largest chromosome in the *O. niloticus* karyotype (Figure 1). This chromosome carries a ZW sex locus in several species of *Oreochromis* [13,46], but not in the *O. niloticus* line studied here. The first ~30Mbp of LG3 shows a standard rate and pattern of recombination. However, the remaining ~70Mbp exhibits almost no recombination in either males or females. This region of highly reduced recombination contains the ZW sex locus in other *Oreochromis* species, and a very high density of repetitive elements.

### Conservation of ancient synteny

Synteny is remarkably conserved among even distantly related teleosts [40,78]. Medaka show few inter-chromosomal rearrangements since shortly after the fish-specific whole genome duplication more than 300 MY ago [34]. Our whole genome alignment of tilapia to medaka supports the previously reported findings that the syntenic organization of teleost genomes is largely stable. The ancestral teleost chromosome number was 24, and contraction of diploid chromosome numbers is usually the result of chromosome fusion and/or translocation events [40]. In cichlids, where the most common chromosome number is 22 [16], we find evidence for two large fusion events on LG7 and LG23 and additional translocations on LG15 and LG17. Cichlid LG7 corresponds to a fusion of medaka chromosomes 6 and 12 (Figure 4), while cichlid LG23 is a fusion of medaka chromosomes 2 and 4 (Figure 5). Clearly, the variation in diploid number observed in other cichlid species implies that there have been additional inter-chromosomal rearrangements, but we predict these will be simple fission/fusion events and not the result of scrambling of these ancient syntenic relationships.

The patterns of recombination across these particular chromosomes provide additional evidence of fusion and translocation events (Figure 4 and Figure 5). There are large deviations from the slope of the recombination curves located precisely where these fusion and translocation events have occurred. This also suggests that the pattern of recombination evolves slowly, as these oddly shaped recombination patterns have persisted for at least 15 million years since the divergence of the common ancestor of *O. niloticus* and the Lake Malawi species. Interestingly, the odd pattern of recombination on LG3 does not seem to be the result of a chromosome fusion event. This lends support to the hypothesis that LG3 accumulated repetitive sequences after it became a sex chromosome, and that this sex chromosome signature and associated recombination suppression persists in *O. niloticus* even following loss of the LG3 sex determination system.

There are many examples of large-scale (>2Mbp) intra-chromosomal rearrangements between *O. niloticus* and Lake Malawi cichlids, as well as rearrangements evident among the Lake Malawi species. In some cases, the anchoring of the *M. zebra* assembly using each map showed the same large structural rearrangement relative to *O. niloticus* for each map (see LG2, LG19, LG20 in Additional File D). This suggests that these rearrangements happened prior to the Lake Malawi radiation, or are specific to *O. niloticus*. In other cases, there are large structural differences relative to *O. niloticus* that are different among the four maps (LG12, Additional File D), which suggests that these rearrangements occurred during the radiation in Lake Malawi. For example, on LG11, the *M. zebra* x *M. mbenjii* map is mostly colinear with *O. niloticus*, but the other three maps show a large rearrangement and some differences in the order of this rearrangement. LG9 of *M. zebra* was particularly difficult to anchor with the *M. mbenjii* x *A. koningsi* map (Table 3). Additional work is needed to better define the structure of these chromosomes in each lineage.

### Evolution of centromere position and sequence

Long-read sequencing has made it possible to assemble centromere repeats [79–81]. A recent study of centromere evolution in medaka provides an example of the role of centromere evolution in speciation [39]. The study showed that the centromere position of many medaka chromosomes has remained unchanged among *Oryzias* species in both acrocentric and non-acrocentric chromosomes. In other chromosomes, the position of centromeres did change and sometimes these chromosomes underwent major structural rearrangements involving other chromosomes. Alignment of the O_niloticus_UMD_NMBU assembly to these new medaka assemblies showed that cichlids have a different set of conserved and variable chromosomes compared to medaka. Additionally, the medaka study showed that centromere sequence repeats were more conserved in the chromosomes that remained acrocentric than in chromosomes that switched between acro- and non-acrocentric or that were non-acrocentric. Assembly and placement of cichlid centromere repeats in multiple species will allow refinement of previous karyotype studies in the context of whole genome assembly comparisons and will also provide insight into centromere evolution at the sequence level. Are there differences in centromere sequence/rate of evolution between acrocentric and non-acrocentric chromosomes? Are these differences great enough to create meiotic incompatibilities in hybrids? Are the positions of centromeres conserved across many species? This study provides a starting point to answer these questions.

### Evolutionary patterns of African cichlid karyotypes

The karyotypes of *O. niloticus* and *M. zebra* in Figure 1 show that there have been at least 5 or 6 changes from subtelo-acrocentric to meta-submetacentric chromosomes. The clearest example of this in the new genome assemblies is the 15Mbp rearrangement on LG23 (Figure 2). Additionally, three similar centromere location changes have occurred, on LG3, LG4 and LG16 (Additional File D). We were able to identify centromere-containing repeats on both the *M. zebra* and *O. niloticus* assemblies in just over half of the chromosomes (LGs 3, 4, 5, 7, 8, 9, 11, 13, 14, 16, 17, 19, 23). The ONSATA (*Oreochromis niloticus* satellite A repeat) and the TZSAT (*Tilapia zillii* satellite repeat) satellite sequences [82] have not been explicitly shown to be centromeric binding sequences, but rather are highly associated with the centromeres via *in situ* hybridization [15]. They are also the only two well-characterized centromere-associated repeat sequences in cichlids. It is possible that these ONSATA and TZSAT repeat sequences may be present in other portions of the chromosome, or that some of them have been assembled incorrectly. Indeed, there are several chromosomes where the ONSATA and TZSAT repeats were identified in multiple distant locations along the chromosome in one or both assemblies (LG6, LG16, LG17, LG19). A centromere was not identified on LG6 in the *M. zebra* assembly, but this chromosome also may have undergone a change in centromere location due to the structural difference between *M. zebra and O. niloticus* (Additional File D).

Two of the chromosomes where we have identified karyotype changes have also been shown to harbor sex-determining loci in African cichlids. The first was the previously mentioned XY sex determination region in *O. niloticus* on LG23 [76]. On LG3, a WZ sex determination region was previously identified [46] and characterized [13] in the congener species *O. aureus*, and now reanalyzed on the O_niloticus_UMD_NMBU assembly (Additional File E). There is a very wide region of sex-patterned differentiation from ~40Mbp to 85Mbp on LG3 in *O. aureus* (13). This region corresponds to the low recombination region in male and female *O. niloticus*. The largest chromosome in the *O. niloticus* karyotype is LG3, and the largest *M. zebra* chromosome is LG7 (Figure 1). Each of these chromosomes is known to be a sex chromosome in their respective lineage. The assembled and anchored chromosomes support these karyotypes since the largest *O. niloticus* assembled chromosome is LG3 and the largest *M. zebra* chromosome is LG7 (Table 1 and Table 3). We suggest that LG3 expanded from the ancestral state in the *O. niloticus* lineage by the accumulation of a large amount of TEs and segmental duplications, likely while linked to sex determination in a basal *Oreochromis* [13]. It is not clear if this apparent runaway elongation of LG3 in *Oreochromis* is due to a sex-determination locus, or if recombination was suppressed first due to some other process. Additional genome assemblies of similar quality in related *Oreochromis* species should allow for further refinement of the evolutionary history of this large sex chromosome in the Oreochromini.

There is also a large (~28Mbp) region of greatly reduced recombination on LG23 in the *O. niloticus* map, as well as in each of the four Lake Malawi maps. LG23 is also the second largest anchored chromosome in the *M. zebra* assembly and third largest chromosome in the *O. niloticus* assembly. It is possible that this arm of LG23 is accumulating TEs similar to LG3, but at an earlier stage. There is an XY sex determination locus on LG23 in *O. niloticus* [57,76], and in at least one species of Lake Victoria cichlid [83], which may be contributing to changes in the size and rate of recombination on this chromosome. Three scenarios may explain these observations: 1) LG23 is an ancient sex chromosome, and though lost in the Malawi lineage, associated recombination suppression remains in Lake Malawi cichlids; 2) The LG23 sex determination locus is indeed segregating in Lake Malawi cichlids but has yet to be identified and described; 3) The recombination pattern on LG23 is not due to sex-chromosome-associated evolution but has been maintained by unknown factors in both lineages.

While many chromosomes have shown extensive rearrangement, it should also be noted that several chromosomes have undergone very little change since the divergence of *M. zebra* and *O. niloticus*. Other than relatively small structural changes at the ends of chromosomes, conserved synteny seems to have been maintained across the entire length of LGs 13, 14, 17 and 18 (Additional File D). It is possible that selective pressures have acted to maintain the synteny of these chromosomes. Since 20% of the *M. zebra* and 10% of the *O. niloticus* genome assemblies remain unanchored, future studies may provide additional structural insights. For example, LG9 in *M. zebra* remains under-anchored. Future *in situ* and physical mapping studies should confirm these results in *O. niloticus* and *M. zebra*. Our work will greatly inform fine-scale cytogenetic studies aimed at characterizing intra-chromosomal differences among cichlid species.

### Recent transposable element expansion in *M. zebra*

Recent evidence has shown that AFC-SINE indels in cis-regulatory regions of genes are highly associated with innovative cichlid phenotypes such as egg-spots [44]. A deletion that may be TE-mediated is responsible for controlling the expression of the *SWS2A* opsin [45]. It is likely that other AFC-specific and other TE-mediated mutations have also contributed to the diverse phenotypes of African cichlids. Therefore, it is important that these TE insertion events are well represented in the genome assemblies.

*M. zebra* has a higher number of recent TE insertions (sequence divergence < 2%) than *O. niloticus* (Figure 6). Since the *O. niloticus* assembly is 43.4Mbp longer than the *M. zebra* assembly, it is possible that the difference in the number of recent TE insertions is even greater than we have quantified here. Each new version of the *M. zebra* genome assembly has increased the total assembled length of many TE super-families (Additional File I), including the AFC-specific families.

We present this finding with several caveats. It is possible that the two species have divergent patterns of insertion across the genome. We previously suggested *O. niloticus* contains larger clusters of repeat arrays that are experiencing recent insertions [13]. These very long arrays do not seem to be present at the same frequency in the *M. zebra* genome. It is possible that many of the recent TE insertions in *O. niloticus* were not assembled and remain hidden in these large arrays. Differences in effective population size (Ne) between the two species may also account for differences in rate of TE accumulation as larger populations will able to purge deleterious insertions more efficiently. Additionally, the DNA of the two species was extracted and sequenced at different times. The *O. niloticus* genome was assembled from 44X coverage of PacBio reads using the P6-C4 chemistry, while the *M. zebra* genome was assembled from 48.5X coverage using the P6-C4 chemistry and 16.5X coverage using a different PacBio chemistry (P5-C3). Other unknown technical factors may also have contributed to the difference that we have described. Future comparisons of additional samples and species assembled using the same sequencing coverage and assembly software/parameters will help to more accurately quantify the recent TE expansion in African Great Lake cichlids.

### Diploid assembly

We present the new *M. zebra* assembly in both haploid and diploid representations. The majority of current genomics tools assume a haploid reference assembly and all subsequent analyses are based on this haploid representation. The use of multiple diploid assemblies will be required to capture population-level patterns of heterozygosity and complex structural variation. The genome assemblies reported here should therefore be considered the beginning of a larger effort to properly represent cichlid genomes. A study of *Arabidopsis thaliana* and *Vitis vinifera* (Cabernet Sauvignon) showed that the phased diploid assemblies produced by FALCON-unzip improved identification of haplotype structure and heterozygous structural variation [7]. Sequencing and assembly of F1 in cattle has also been shown to recover these complex regions better and may be the way forward for assembly of diploid genomes [84]. Graph genome representations [85,86] have been shown to improve variant calling in complex regions such as the human leukocyte antigen (HLA) [87], major histocompatibility complex (MHC) [88] and centromeres [89]. Additional long-read diploid assemblies will be able to better represent genetic variation, particularly in regions of complex variation which current long read assemblies are beginning to span [80].

## Potential implications

This study highlights the evolutionary insights that can be gained using a comparison of high-quality chromosome-scale genome assemblies, genetic recombination maps and cytogenetics across multiple related and, in this case, rapidly evolving species. It further illustrates the need for high-quality, chromosome-scale genome assemblies for answering many basic biological questions. This study illustrates the structural changes that can occur in the genomes of a rapidly evolving clade. It will be interesting to make comparisons to other radiations in the tree of life, both large and small. This study provides a wide-angle view of African cichlid genome history and demonstrates how these high-quality resources can be used for many different types of evolutionary genomic analyses. As additional high-quality cichlid genomes are generated, this study will provide the foundation for comparisons of structural variation, recombination, cytogenetics, and repetitive sequences across the cichlid phylogeny. Many new questions have been generated here. How do the structural changes of African cichlid genomes compare to other groups? Is the pattern of few inter-chromosomal, but many intra-chromosomal differences seen here found in additional Lake Malawi genera as well as other radiations in Lake Tanganyika and Lake Victoria? Are these patterns of recombination observed across the majority of cichlids? Are any deviations from these typical recombination patterns related to specific phenotypic traits or sex chromosome history? How have these chromosomes evolved structurally? We look forward to the new dawn in cichlid genomics.

## Methods

### *O. niloticus* SNP array map, misassembly detection and new anchoring

Offspring (n=689) and parents from 41 full-sib families belonging to the 20^th^, 24^th^ and 25^th^ generations of the GST^®^ strain were analyzed using a custom 57K SNP Axiom^®^ Nile Tilapia Genotyping Array [22]. SNPs classified as “PolyHighRes” or “No-MinorHom” by Axiom Analysis Suite (Affymetrix, Santa Clara, USA), and having a minor-allele frequency ≥ 0.05, and call rate ≥ 0.85 were used in genetic map construction (n= 40,548). Lep-MAP2 [90] was used to order these SNPs into linkage groups in a stepwise process beginning with SNPs being assigned to linkage groups using the ‘SeparateChromosomes’ command. LOD thresholds were adjusted until 22 linkage groups, which correspond with the *O. niloticus* karyotype. Unassigned SNPs were subsequently added to linkage groups using the ‘JoinSingles’ command and a more relaxed LOD threshold, and ordered within each linkage group using the ‘OrderMarkers’ command.

Sequence flanking each SNP (2 x 35nt) was used to precisely position 40,190 SNPs to the O_niloticus_UMD1 assembly (NCBI accession MKQE00000000) and thereby integrate the linkage and physical maps. This revealed 22 additional contig misassemblies (i.e. contigs containing SNPs from different LGs) that were not detected in the original anchoring for O_niloticus_UMD1. These contigs that were subsequently broken. Linkage information was subsequently used to order and orientate contigs and build sequences for 22 Nile tilapia LGs in the new O_niloticus_UMD_NMBU assembly following the previous cichlid nomenclature [5,13,56,91].

LD results (r^2^ > 0.97) presented in Figure 4, Figure. 5 and Additional file F, were produced in PLINK2 version 1.90b3w [92] using the pedigree described above and SNP-positions given in [92].

### PacBio Sequencing of *M. zebra*

The previous version of the *M. zebra* assembly, M_zebra_UMD1 [12], included 16.5X PacBio sequencing (25 SMRT cells using the P5-C3 chemistry) on an PacBio RS II machine [12]. An additional library was prepared using the same Qiagen MagAttract HMW DNA extraction and Blue Pippin pulse-field gel electrophoresis size selection. An additional 60 SMRT cells (using the P6-C4 chemistry) were sequenced on the same PacBio RS II at the University of Maryland Genomics Resource Center as the previous 16.5X P5-C3 data. These P6-C4 SMRT cells comprised ~48.5X coverage to bring combined total to ~65X coverage.

### *M. zebra* diploid genome assembly

The 65X PacBio reads were assembled using FALCON-integrate/FALCON_unzip (*version 0.4.0*) [7]. The following parameters were used for the ‘*fc_run.py*’ assembly step:

> *length_cutoff = 9000*
>
> *length_cutoff_pr = 9000*
>
> *pa_HPCdaligner_option = -v -dal128 -H10000 -M60 -t16 -e.70 -l2000 -s100 -k14 -h480 -w8*
>
> *ovlp_HPCdaligner_option = -v -dal128 -H10000 -M60 -t32 -h1024 -e.96 -l1000 -s100 -n k24*
>
> *falcon_sense_option = --output_multi --min_idt 0.70 --min_cov 4 --max_n_read 350 -- n_core 5*
>
> *overlap_filtering_setting = --max_diff 100 --max_cov 150 --min_cov 0 --bestn 10 -- n_core 18*

This was followed by the unzip step (‘*fc_unzip.py*’) and quiver polishing of the diploid assembly with the ‘*fc_quiver.py*’ assembly step.

### Polishing of the *M. zebra* diploid genome assembly

The diploid assembly described above includes a PacBio polishing (quiver) step. However, there were also Illumina reads available for *M. zebra* from the first version of the assembly [5]. Trimming and filtering of the raw *M. zebra* Illumina reads are described for the previous version of the assembly [12]. The trimmed and filtered fragment library corresponded to 30.1X coverage and the trimmed and filtered 2-3kb library corresponded to 32.6X coverage for a total of 62.7X Illumina coverage. These Illumina reads were aligned to the diploid assembly with BWA mem [93] (*version 0.7.12-r1044*). Pilon [94] (*version 1.22*) was run supplying the fragment library with the ‘--*frags*' option, the 2-3kb library with the ‘--*jumps*’ option and the following options: ‘--*diploid*--*fix bases* --*mindepth 10* --*minmq 1* --*minqual 1 --nostrays*’.

This intermediate, Illumina-polished assembly was then polished again with the PacBio reads using SMRT-Analysis [95] (*version 2.3.0.140936*) using the 65X raw PacBio reads. First, each SMRT cell was separately aligned to the intermediate polished assembly using pbalign (*version 0.2.0.138342*) with the ‘--*forQuiver*' flag. Next, cmph5tools.py (*version 0.8.0*) was used to merge and sort (with the ‘--*deep*’ flag) the pbalign .h5 output files for each SMRT cell. Finally, Quiver (*GenomicConsensus version 0.9.2* and *ConsensusCore version 0.8.8*) was run on the merged and sorted pbalign output to produce an initial polished assembly.

### Detecting misassemblies in *M. zebra*

To detect misassemblies present in the intermediate polished assembly, several datasets were analyzed and compared. This included four genetic maps: A genetic map with 834 markers generated from RAD genotyping of 160 F2 individuals from a cross of *M. zebra* and *M. mbenjii* [49]; a genetic map with 946 markers generated from RAD genotyping of 262 F2 individuals from a cross of *Labeotropheus fuelleborni* and *Tropheops* ‘*red cheek*’ [50]; a genetic map of 2,553 markers generated from RAD genotyping of 331 F2 individuals from a cross of *M. mbenjii* and *Aulonocara koningsi* (cross and map construction details in separate Methods section); a genetic map of 1,217 markers generated from RAD genotyping of 161 F2 individuals from a cross of *M. mbenjii* and *A. baenschi* (cross and map construction details in separate Methods section).

The markers for each of the four maps were aligned to the intermediate polished assembly using BWA mem [93] (*version 0.7.12-r1044*) and a separate SAM file was generated. Chromonomer [96] (*version 1.05*) was run for each map using these respective SAM files and map information as input. Chromonomer detected contigs in the intermediate assembly that were mapped to multiple linkage groups.

To narrow the location of these identified misassemblies, the Illumina 40kb mate-pair library from the first *M. zebra* assembly [5] was aligned to the intermediate assembly. The raw PacBio reads were aligned using BLASR [97] (version 1.3.1.127046) with the following parameters: ‘*-minMatch 8 -minPctI-dentity 70 -bestn 1 -nCandidates 10 -maxScore -500 -nproc 40 -noSplitSubreads –sam*’ Regions of abnormal coverage in the PacBio read alignments as well as abnormal clone coverage in the 40kb mate-pair were identified for most potential misassemblies identified by the genetic maps. These misassembly regions were manually inspected using these alignments in IGV [98]. Additionally, RefSeq [51] (*release 76*) *M. zebra* transcripts were aligned to the intermediate assembly using GMAP [99] (*version 2015-07-23*) and RepeatMasker [100] repeat annotations were considered when defining the exact location of a misassembly break.

One additional misassembly was identified during the comparison of linkage maps (next section) and was subsequently broken using the same process as above.

### *M. zebra* assembly anchoring

The same four genetics maps used above for misassembly detection were also used for anchoring the assembly contigs (after breaking) into the final set of linkage groups. Chromonomer [96] (*version 1.05*) was run on each of these four genetic maps to anchor the polished and misassembly corrected contigs. BWA mem (*version 0.7.12-r1044*) was used to create the input SAM file by aligning each respective map marker sequences to these contigs. Gaps of 100bp were placed between anchored contigs. The final M_zebra_UMD2 anchoring was generated by anchoring LG9 and LG11 with the *M. zebra* and *M. mbenjii* map [49], LG20 with the *M. mbenjii* and *A. baenschi* map and the remaining 19 LGs with the *M. mbenjii* and *A. koningsi* map. To accomplish this anchoring, the markers for each of those respective maps and LGs were used with Chromonomer as described above.

### *M. zebra* repeat annotation

RepeatModeler [101] (*version open-1.0.8*) was first used to identify and classify *de novo* repeat families present in the final anchored assembly. These *de novo* repeats were combined with the RepBase-derived RepeatMasker libraries [102]. RepeatMasker [100] (*version open-4.0.5*) was run on the final anchored assembly using NCBI BLAST+ (*version 2.3.0+*) as the engine (‘-*e ncbi*’) and specifying the combined repeat library (‘-*lib*’). The more sensitive slow search mode (‘-s’) was used. The repeat landscape was generated with the RepeatMasker ‘*calcDivergenceFromAlign.pl*’ and ‘*createRepeatLandscape.pl*’ utility scripts.

### *M. zebra* BUSCO genome-completeness analysis

BUSCO (*version 3.0.2*) was run on the M_zebra_UMD2 anchored assembly in the genome mode (-*m geno*) and compared against the vertebrate BUSCO set (‘vertebrata_odb9’).

### Whole genome alignment of *M. zebra* to *O. niloticus*

The final anchored M_zebra_UMD2 assembly was aligned to the O_niloticus_UMD_NMBU assembly using the *‘nucmer’* program of the MUMmer package [103] (*version 3.1*). The default *nucmer* parameters were used and the raw *nucmer* alignments were filtered using the ‘*delta-filte*’ program with the following options: ‘-*o 50 -l 50 -1 -i 10 -u 10*’. These filtered alignments were converted to a tab-delimited set of coordinates using the ‘*show-coords*’ program with the following options: ‘-*I 10 -L 5000 -l -T-H*'. This set of coordinates was then visualized using Ribbon [104] and used to generate the images in Additional File D.

### Whole genome alignment of *M. zebra* to medaka

The HSOK medaka genome assembly version 2.2.4 was downloaded from http://utgenome.org/medaka_v2/#!Assembly.md and corresponds to NCBI accession (GCA_002234695.1). Similar to the M_zebra_UMD2 comparison, O_niloticus_UMD_NMBU was aligned to the medaka HSOK genome with *nucmer*. The ‘*delta-filter*’ settings were adjusted to *‘-1 -l 50 -i 50 -u 50*’ to account for the increased divergence between the two more distantly related species. The ‘*show-coords’* settings were also adjusted to ‘-*I 50 -L 50 -l -T -H*’. Alignments were again viewed with Ribbon to identify putative chromosome fusion and translocation events and used to generate the part of the images in Figure 4 and Figure 5.

## Declarations

**List of abbreviations**
AFC –: African cichlid specific repetitive element.
BUSCO –: Benchmarking Universal Single-Copy Orthologs.
cM –: Centimorgan.
LD –: Linkage disequilibrium.
LG –: Linkage group.
LINE –: Long interspersed nuclear element.
LGs –: Linkage groups.
N50 –: Shortest contig/scaffold/read/sequence length at 50% of the genome/read set.
NG50 –: Shortest contig/scaffold/read/sequence length at 50% of the estimated genome/read set size.
ONSATA –: *Oreochromis niloticus* satellite A repeat.
ONSATB –: *Oreochromis niloticus* satellite B repeat PacBio – Pacific Biosciences.
RAD –: Restriction site associated DNA.
RefSeq –: NCBI Reference Sequence Database.
SMRT –: Single Molecule, Real-Time.
TE –: Transposable element.
TZSAT –: *Tilapia zillii* satellite repeat.

## Consent for publication

Not applicable.

## Competing interests

The authors declare that they have no competing interests.

## Funding

This work was supported by the United States Department of Agriculture under project number MD.W-2014-05906 to TDK, the National Science Foundation under grant number DEB-1143920 to TDK, the National Institutes of Health project R01-EY024639 to KLC and the Beckman Young Investigator Award from the Arnold and Mabel Beckman Foundation to RBR.

## Author's contributions

MAC, TDK, and KLC conceived the study. TDK carried out HMW DNA extraction. MAC carried out computational analyses. RJ, ECM, SPN, RBR and SL performed genetic map construction. MAC and SL integrated the tilapia linkage map with the assembly. WJG organized map data for anchoring. MAC and TDK wrote the manuscript. All authors read and approved the manuscript.

## Acknowledgements

We acknowledge all members of the UMD cichlid labs for their countless discussions on this project. We thank Luke Tallon and Naomi Sengamalay at the Genomics Resource Center, Institute for Genome Sciences for providing a high quality PacBio library and sequence reads. We acknowledge the University of Maryland supercomputing resources (www.it.umd.edu/hpcc/) made available in conducting the research reported in this paper. We thank Silje Karoliussen and Matthew Kent for genotyping and quality filtering of the Nile tilapia 58K SNP-array data. We thank many individuals in the cichlid community for their patience while we developed these resources.

## Author’s information

Not applicable.

## Availability of supporting data and materials

The *O. niloticus* Whole Genome Shotgun project has been deposited at DDBJ/ENA/GenBank under the accession MKQE00000000 (O_niloticus_UMD1). The version described in this paper is version MKQE02000000 (O_niloticus_UMD_NMBU). The *M. zebra* Whole Genome Shotgun project has been deposited at DDBJ/ENA/GenBank under the accession AGTA00000000. The version described in this paper is version AGTA05000000.

## Endnotes

Not applicable.

## Additional files

Additional File A - Size distribution of the M_zebra_UMD2 assembled haplotigs and theoretical recombination rate for several different effective population sizes.

Additional File B – M_zebra_UMD2 FALCON p-contigs where markers from two or more different LGs maps aligned, indicating a potential inter-LG misassembly.

Additional File C – Screenshot of IGV view to inspect potential misassemblies. In this example, a misassembly on this contig was confirmed at position 420,665 (indicated by the white arrows). The top red box shows the portion of the contig that is being visualized. LG17 markers aligned at 186kb and 308kb, while LG10a markers aligned at 760kb and 1.6Mbp as indicated by the red arrows. The top two tracks below that are the read coverage plots for the PacBio read alignments against the diploid and haploid sets of contigs. There is a sharp decrease in PacBio read coverage at the misassembly location. The track below that show 40kb mate-pair alignments and also show no clone coverage at the location of the misassembly.

Additional File D – *M. zebra* assembly contigs anchored with each of the 4 maps and aligned to O_niloticus_UMD_NMBU (indicated as black on bottom with contigs in red for each panel). Centromeres indicated with black triangles. Contigs are represented as red lines above each respective assembly.

Additional File E – (a) F_ST_ comparison of male and female *O. aureus* LG3 WZ. (b) *O. niloticus* recombination curve of LG3 from Additional File F.

Additional File F – *O. niloticus* recombination curves for females (red) and males (blue). Centromere repeats are displayed as green triangles where applicable. X-axis represent the location along the anchored LG. Left Y-axis represents linkage disequilibrium (black points, r^2^ > 0.97) and right Y-axis shows the map location for each marker.

Additional File G – Comparison of recombination in the four Lake Malawi genetic maps. LGs from maps that needed to be reversed from their original published order are indicated in the legend. The detected misassembly is included as “LG12 misassembly” on page 13.

Additional File H – Comparison of the repeat landscape in the *M. zebra* and *O. niloticus* genome assemblies using same assembly parameters.

Additional File I – Comparison of the repeat landscape in the three *M. zebra* assembly versions.

Additional File J – Table of the orientation of Lake Malawi recombination maps for each LG. The forward and reverse orientation information of each map was used to generate recombination plots in the same orientation for Additional File F.

## References

1. Kocher TD. Adaptive evolution and explosive speciation: the cichlid fish model. Nat Rev Genet. 2004;5:288–98.

2. Malinsky M, Challis RJ, Tyers a M, Schiffels S, Terai Y, Ngatunga BP, et al. Genomic islands of speciation separate cichlid ecomorphs in an East African crater lake. Science (80-). 2015;350:1493–8.

3. Malinsky M, Svardal H, Tyers AM, Miska EA, Genner MJ, Turner GF, et al. Whole Genome Sequences Of Malawi Cichlids Reveal Multiple Radiations Interconnected By Gene Flow. bioRxiv. 2017;143859.

4. Meier JI, Sousa VC, Marques DA, Selz OM, Wagner CE, Excoffier L, et al. Demographic modelling with whole-genome data reveals parallel origin of similar Pundamilia cichlid species after hybridization. Mol Ecol. 2017;26:123–41.

5. Brawand D, Wagner CE, Li YI, Malinsky M, Keller I, Fan S, et al. The genomic substrate for adaptive radiation in African cichlid fish. Nature. 2014;513:375–81.

6. Bradnam KR, Fass JN, Alexandrov A, Baranay P, Bechner M, Birol İ, et al. Assemblathon 2: evaluating de novo methods of genome assembly in three vertebrate species. Gigascience. 2013;2.

7. Chin C-S, Peluso P, Sedlazeck FJ, Nattestad M, Concepcion GT, Clum A, et al. Phased Diploid Genome Assembly with Single Molecule Real-Time Sequencing. Nat Methods. 2016;13:1050–4.

8. Koren S, Walenz BP, Berlin K, Miller JR, Bergman NH, Phillippy AM. Canu: Scalable and accurate long-read assembly via adaptive κ-mer weighting and repeat separation. Genome Res. 2017;27:722–36.

9. Chaisson MJP, Huddleston J, Dennis MY, Sudmant PH, Malig M, Hormozdiari F, et al. Resolving the complexity of the human genome using single-molecule sequencing. Nature. Nature Publishing Group; 2014;517:608–11.

10. Gordon D, Huddleston J, Chaisson MJP, Hill CM, Kronenberg ZN, Munson KM, et al. Long-read sequence assembly of the Gorilla Genome. Science (80-). 2016;352.

11. Zimin A V., Puiu D, Hall R, Kingan S, Clavijo BJ, Salzberg SL. The first near-complete assembly of the hexaploid bread wheat genome, Triticum aestivum. Gigascience. 2017;

12. Conte MA, Kocher TD. An improved genome reference for the African cichlid, Metriaclima zebra. BMC Genomics. BMC Genomics; 2015;16:724.

13. Conte MA, Gammerdinger WJ, Bartie KL, Penman DJ, Kocher TD. A high quality assembly of the Nile Tilapia (Oreochromis niloticus) genome reveals the structure of two sex determination regions. BMC Genomics. BMC Genomics; 2017;18:341.

14. Feldberg E, Ivan J, Porto R, Antonio L, Bertollo C. Chromosomal Changes and Adaptation Of Cichlid Fishes During Evolution. AL Val BG Kapoor (eds), Fish Adapt. 2003;285–309.

15. Ferreira IA, Poletto AB, Kocher TD, Mota-Velasco JC, Penman DJ, Martins C. Chromosome evolution in African cichlid fish: contributions from the physical mapping of repeated DNAs. Cytogenet Genome Res. 2010;129:314–22.

16. Poletto AB, Ferreira IA, Cabral-de-Mello DC, Nakajima RT, Mazzuchelli J, Ribeiro HB, et al. Chromosome differentiation patterns during cichlid fish evolution. BMC Genet. 2010;11:50.

17. Carbone L, Alan Harris R, Gnerre S, Veeramah KR, Lorente-Galdos B, Huddleston J, et al. Gibbon genome and the fast karyotype evolution of small apes. Nature. 2014;513:195–201.

18. Damas J, O’Connor R, Farré M, Lenis VPE, Martell HJ, Mandawala A, et al. Upgrading short-read animal genome assemblies to chromosome level using comparative genomics and a universal probe set. Genome Res. 2017;27:875–84.

19. Lewin HA, Larkin DM, Pontius J, O’Brien SJ. Every genome sequence needs a good map. Genome Res. 2009;19:1925–8.

20. Hellmann I, Ebersberger I, Ptak SE, Pääbo S, Przeworski M. A Neutral Explanation for the Correlation of Diversity with Recombination Rates in Humans. Am J Hum Genet. 2003;72:1527–35.

21. Wolf JBW, Ellegren H. Making sense of genomic islands of differentiation in light of speciation. Nat Rev Genet. Nature Publishing Group; 2017;18:87–100.

22. Joshi R, Tola A, Kent M, Sciences A, Joshi R. Development and validation of 58K SNP-array and high-density linkage map in Nile tilapia (*O. niloticus*). bioRxiv. 2018;

23. Begun DJ, Aquadro CF. Levels of naturally occurring DNA polymorphism correlate with recombination rates in D. melanogaster. Nature. 1992;356:519–20.

24. Kulathinal RJ, Bennett SM, Fitzpatrick CL, Noor MAF. Fine-scale mapping of recombination rate in Drosophila refines its correlation to diversity and divergence. Proc Natl Acad Sci. 2008;105:10051–6.

25. Charlesworth D. Evolution of recombination rates between sex chromosomes. Phil Trans R Soc B. 2017;372:20160456.

26. Stapley J, Feulner PGD, Johnston SE, Santure AW, Smadja CM. Variation in recombination frequency and distribution across eukaryotes: Patterns and processes. Philos Trans R Soc B Biol Sci. 2017;

27. Gante HF, Matschiner M, Malmstrøm M, Jakobsen KS, Jentoft S, Salzburger W. Genomics of speciation and introgression in Princess cichlid fishes from Lake Tanganyika. Mol Ecol. 2016;

28. Werren JH. Selfish genetic elements, genetic conflict, and evolutionary innovation. Proc Natl Acad Sci U S A. 2011;108 Suppl:10863–70.

29. Ser JR, Roberts RB, Kocher TD. Multiple interacting loci control sex determination in Lake Malawi cichlid fish. Evolution (N Y). 2010;64:486–501.

30. Roberts RB, Ser JR, Kocher TD. Sexual conflict resolved by invasion of a novel sex determiner in Lake Malawi cichlid fishes. Science. 2009;326:998–1001.

31. Gammerdinger WJ, Conte MA, Acquah EA, Roberts RB, Kocher TD. Structure and decay of a proto-Y region in Tilapia, Oreochromis niloticus. BMC Genomics. 2014;15:975.

32. Gammerdinger WJ, Conte MA, Baroiller J-F, D’Cotta H, Kocher TD. Comparative analysis of a sex chromosome from the blackchin tilapia, Sarotherodon melanotheron. BMC Genomics. 2016;17:808.

33. Clark FE, Conte MA, Ferreira-Bravo IA, Poletto AB, Martins C, Kocher TD. Dynamic sequence evolution of a sex-Associated b chromosome in lake Malawi cichlid fish. J Hered. 2017;108:53–62.

34. Kasahara M, Naruse K, Sasaki S, Nakatani Y, Qu W, Ahsan B, et al. The medaka draft genome and insights into vertebrate genome evolution. Nature. 2007;447:714–9.

35. Roberts NB, Juntti SA, Coyle KP, Dumont BL, Stanley MK, Ryan AQ, et al. Polygenic sex determination in the cichlid fish. BMC Genomics. 2016;1–13.

36. Mazzuchelli J, Kocher TD, Yang F, Martins C. Integrating cytogenetics and genomics in comparative evolutionary studies of cichlid fish. BMC Genomics. 2012;13.

37. Liu F, Sun F, Li J, Xia JH, Lin G, Tu RJ, et al. A microsatellite-based linkage map of salt tolerant tilapia (*Oreochromis mossambicus* x *Oreochromis spp*.) and mapping of sex-determining loci. BMC Genomics. 2013;14:1–14.

38. Gammerdinger WJ, Conte MA, Baroiller J, Cotta HD, Kocher TD. Comparative analysis of a sex chromosome from the blackchin tilapia, Sarotherodon melanotheron. BMC Genomics. 2016;1–10.

39. Ichikawa K, Tomioka S, Suzuki Y, Nakamura R, Doi K, Yoshimura J, et al. Centromere evolution and CpG methylation during vertebrate speciation. Nat Commun. 2017;8:1833.

40. Amores A, Catchen J, Nanda I, Warren W, Walter R, Schartl M, et al. A RAD-tag genetic map for the platyfish (Xiphophorus maculatus) reveals mechanisms of karyotype evolution among teleost fish. Genetics. 2014;197:625–41.

41. Takahashi K, Terai Y, Nishida M, Okada N. A Novel Family of Short Interspersed Repetitive Elements (SINEs) from Cichlids : The Patterns of Insertion of SINEs at Orthologous Loci Support the Proposed Monophyly of Four Major Groups of Cichlid Fishes in Lake Tanganyika. Mol Biol Evol. 1998;15:391–407.

42. Takahashi K, Okada N. Mosaic structure and retropositional dynamics during evolution of subfamilies of short interspersed elements in African cichlids. Mol Biol Evol. 2002;19:1303–12.

43. Chuong EB, Elde NC, Feschotte C. Regulatory activities of transposable elements: From conflicts to benefits. Nat Rev Genet. 2017;18:71–86.

44. Santos ME, Braasch I, Boileau N, Meyer BS, Sauteur L, Böhne A, et al. The evolution of cichlid fish egg-spots is linked with a cis-regulatory change. Nat Commun. 2014;5:5149.

45. Schulte JE, O’Brien CS, Conte M a, O’Quin KE, Carleton KL. Interspecific variation in Rx1 expression controls opsin expression and causes visual system diversity in African cichlid fishes. Mol Biol Evol. 2014;31:2297–308.

46. Lee B, Hulata G, Kocher TD. Two unlinked loci controlling the sex of blue tilapia (*Oreochromis aureus*). Heredity (Edinb). 2004;92:543–9.

47. Gregory TR. Animal Genome Size Database [Internet]. 2016. Available from: http://www.genomesize.com

48. Husemann M, Nguyen R, Ding B, Danley PD. A genetic demographic analysis of Lake Malawi rock-dwelling cichlids using spatio-temporal sampling. Mol Ecol. 2015;24:2686–701.

49. O’Quin CT, Drilea AC, Conte MA, Kocher TD. Mapping of pigmentation QTL on an anchored genome assembly of the cichlid fish, Metriaclima zebra. BMC Genomics. 2013;14:1.

50. Albertson RC, Powder KE, Hu Y, Coyle KP, Roberts RB, Parsons KJ, et al. Genetic basis of continuous variation in the levels and modular inheritance of pigmentation in cichlid fishes. Mol Ecol. 2014;23:5135–50.

51. O’Leary NA, Wright MW, Brister JR, Ciufo S, Haddad D, McVeigh R, et al. Reference sequence (RefSeq) database at NCBI: Current status, taxonomic expansion, and functional annotation. Nucleic Acids Res. 2016;44:D733–45.

52. Franck JPC, Kornfield I, Wright JM. The utility of sata satellite dna sequences for inferring phylogenetic relationships among the three major genera of tilapiine cichlid fishes. Mol Phylogenet Evol. 1994;3:10–6.

53. Melters DP, Bradnam KR, Young H a, Telis N, May MR, Ruby JG, et al. Comparative analysis of tandem repeats from hundreds of species reveals unique insights into centromere evolution. Genome Biol. BioMed Central Ltd; 2013;14:R10.

54. Kumar S, Stecher G, Suleski M, Hedges SB. TimeTree: A Resource for Timelines, Timetrees, and Divergence Times. Mol Biol Evol. 2017;34:1812–9.

55. Ferreira IA, Martins C. Physical chromosome mapping of repetitive DNA sequences in Nile tilapia *Oreochromis niloticus*: Evidences for a differential distribution of repetitive elements in the sex chromosomes. Micron. 2008;39:411–8.

56. Guyon R, Rakotomanga M, Azzouzi N, Coutanceau JP, Bonillo C, D’Cotta H, et al. A high-resolution map of the Nile tilapia genome: a resource for studying cichlids and other percomorphs. BMC Genomics. 2012;13:222.

57. Eshel O, Shirak A, Weller JI, Hulata G, Ron M. Linkage and Physical Mapping of Sex Region on LG23 of Nile Tilapia (*Oreochromis niloticus*). G3 Genes, Genomes, Genet. 2012;2:35–42.

58. Peterson EN, Cline ME, Moore EC, Roberts NB, Roberts RB, Moore EC. Genetic sex determination in *Astatotilapia calliptera*, a prototype species for the Lake Malawi cichlid radiation. The Science of Nature; 2017;

59. Terai Y, Takahashi K, Okada N. SINE cousins: the 3’-end tails of the two oldest and distantly related families of SINEs are descended from the 3’ ends of LINEs with the same genealogical origin. Mol Biol Evol. 1998;15:1460–71.

60. Simão FA, Waterhouse RM, Ioannidis P, Kriventseva E V., Zdobnov EM. BUSCO: Assessing genome assembly and annotation completeness with single-copy orthologs. Bioinformatics. 2015;31:3210–2.

61. Waterhouse RM, Seppey M, Simao FA, Manni M, Ioannidis P, Klioutchnikov G, et al. BUSCO applications from quality assessments to gene prediction and phylogenomics. Mol Biol Evol. 2018;

62. Maylandia zebra Annotation Report [Internet]. [cited 2018 May 23]. Available from: https://www.ncbi.nlm.nih.gov/genome/annotation_euk/Maylandia_zebra/103/

63. Maylandia zebra Annotation Report [Internet]. [cited 2018 May 23]. Available from: https://www.ncbi.nlm.nih.gov/genome/annotation_euk/Maylandia_zebra/104/

64. Dolgin ES, Charlesworth B. The effects of recombination rate on the distribution and abundance of transposable elements. Genetics. 2008;178:2169–77.

65. Sakamoto T, Danzmann RG, Gharbi K, Howard P, Ozaki A, Khoo SK, et al. A microsatellite linkage map of rainbow trout (*Oncorhynchus mykiss*) characterized by large sex-specific differences in recombination rates. Genetics. 2000;155:1331–45.

66. Moen T, Hoyheim B, Munck H, Gomez-Raya L. A linkage map of Atlantic salmon (*Salmo salar*) reveals an uncommonly large difference in recombination rate between the sexes. Anim Genet. 2004;35:81–92.

67. Gharbi K, Gautier A, Danzmann RG, Gharbi S, Sakamoto T, Høyheim B, et al. A linkage map for brown trout (*Salmo trutta*): Chromosome homeologies and comparative genome organization with other salmonid fish. Genetics. 2006;172:2405–19.

68. Rexroad CE, Palti Y, Gahr SA, Vallejo RL. A second generation genetic map for rainbow trout (*Oncorhynchus mykiss*). BMC Genet. 2008;9:1–14.

69. Sardell JM, Cheng C, Dagilis AJ, Ishikawa A, Kitano J, Peichel CL, et al. Sex Differences in Recombination in Sticklebacks. G3 Genes, Genomes, Genet. 2018;

70. Castaño-Sánchez C, Fuji K, Ozaki A, Hasegawa O, Sakamoto T, Morishima K, et al. A second generation genetic linkage map of Japanese flounder (*Paralichthys olivaceus*). BMC Genomics. 2010;11.

71. Singer A, Perlman H, Yan Y, Walker C, Corley-Smith G, Brandhorst B, et al. Sex-specific recombination rates in zebrafish (*Danio rerio*). Evol Heal Dis. 2002;160:649–57.

72. Roesti M, Moser D, Berner D. Recombination in the threespine stickleback genome - Patterns and consequences. Mol Ecol. 2013;22:3014–27.

73. Zeng Q, Fu Q, Li Y, Waldbieser G, Bosworth B, Liu S, et al. Development of a 690 K SNP array in catfish and its application for genetic mapping and validation of the reference genome sequence. Sci Rep. 2017;7:1–14.

74. Charlesworth B. The evolution of sex chromosomes. Science (80-). 1991;251:1030–3.

75. Cnaani A, Lee BY, Zilberman N, Ozouf-Costaz C, Hulata G, Ron M, et al. Genetics of sex determination in tilapiine species. Sex Dev. 2008;2:43–54.

76. Li M, Sun Y, Zhao J, Shi H, Zeng S, Ye K, et al. A Tandem Duplicate of Anti-Müllerian Hormone with a Missense SNP on the Y Chromosome Is Essential for Male Sex Determination in Nile Tilapia, *Oreochromis niloticus*. PLoS Genet. 2015;11:1–23.

77. O’Quin CT. Thesis Dissertation [Internet]. University of Maryland; 2014. Available from: http://hdl.handle.net/1903/15212

78. Guyomard R, Boussaha M, Krieg F, Hervet C, Quillet E. A synthetic rainbow trout linkage map provides new insights into the salmonid whole genome duplication and the conservation of synteny among teleosts. BMC Genet. 2012;13:1–12.

79. Sevim V, Bashir A, Chin CS, Miga KH. Alpha-CENTAURI: Assessing novel centromeric repeat sequence variation with long read sequencing. Bioinformatics. 2016;32:1921–4.

80. Jain M, Koren S, Quick J, Rand AC, Sasani TA, Tyson JR, et al. Nanopore sequencing and assembly of a human genome with ultra-long reads. Nat Biotechnol. 2018;

81. Jain M, Olsen HE, Turner DJ, Stoddart D, Bulazel K V., Paten B, et al. Linear Assembly of a Human Y Centromere using Nanopore Long Reads. bioRxiv. 2017;

82. Franck J, Wright J, McAndrew B. Genetic variability in a family of satellite DNAs from tilapia (Pisces: Cichlidae). Genome. 1992;35:719–25.

83. Feulner PGD, Schwarzer J, Haesler MP, Meier JI, Seehausen O. A dense linkage map of Lake Victoria cichlids improved the Pundamilia genome assembly and revealed a major QTL for sex-determination. G3 Genes, Genomes, Genet. 2018;8:2411–20.

84. Koren S, Rhie A, Walenz BP, Dilthey AT, Bickhart DM, Kingan SB, et al. Complete assembly of parental haplotypes with trio binning. bioRxiv. 2018;271486.

85. Novak AM, Hickey G, Garrison E, Blum S, Connelly A, Dilthey A, et al. Genome Graphs. bioRxiv. 2017;101378.

86. Paten B, Novak AM, Eizenga JM, Garrison E. Genome graphs and the evolution of genome inference. Genome Res. 2017;27:665–76.

87. Eggertsson HP, Jonsson H, Kristmundsdottir S, Hjartarson E, Kehr B, Masson G, et al. Graphtyper enables population-scale genotyping using pangenome graphs. Nat Genet. 2017;49:1654–60.

88. Dilthey A, Cox C, Iqbal Z, Nelson MR, McVean G. Improved genome inference in the MHC using a population reference graph. Nat Genet. 2015;47:682–8.

89. Miga KH, Newton Y, Jain M, Altemose N, Willard HF, Kent WJ. Centromere reference models for human chromosomes X and Y satellite arrays. 2014;697–707.

90. Rastas P, Calboli FCF, Guo B, Shikano T, Merilä J. Construction of Ultradense Linkage Maps with Lep-MAP2: Stickleback F2 Recombinant Crosses as an Example. Genome Biol Evol. 2015;8:78–93.

91. Lee B-Y, Lee W-J, Streelman JT, Carleton KL, Howe AE, Hulata G, et al. A second-generation genetic linkage map of tilapia (*Oreochromis spp*.). Genetics. 2005;170:237–44.

92. Chang CC, Chow CC, Tellier LCAM, Vattikuti S, Purcell SM, Lee JJ. Second-generation PLINK: Rising to the challenge of larger and richer datasets. Gigascience. 2015;4.

93. Li H. Aligning sequence reads, clone sequences and assembly contigs with BWA-MEM. arXiv. 2013;00:1–2.

94. Walker BJ, Abeel T, Shea T, Priest M, Abouelliel A, Sakthikumar S, et al. Pilon: An integrated tool for comprehensive microbial variant detection and genome assembly improvement. PLoS One. 2014;9.

95. PacificBiosciences/SMRT-Analysis [Internet]. [cited 2014 May 5]. Available from: https://github.com/PacificBiosciences/SMRT-Analysis

96. Catchen J, Amores A. Chromonomer [Internet]. Available from: http://catchenlab.life.illinois.edu/chromonomer/

97. Chaisson MJ, Tesler G. Mapping single molecule sequencing reads using basic local alignment with successive refinement (BLASR): application and theory. BMC Bioinformatics. 2012;13.

98. Thorvaldsdóttir H, Robinson JT, Mesirov JP. Integrative Genomics Viewer (IGV): High-performance genomics data visualization and exploration. Brief Bioinform. 2013;14:178–92.

99. Wu TD, Watanabe CK. GMAP: a genomic mapping and alignment program for mRNA and EST sequences. Bioinformatics. 2005;21:1859–75.

100. Smit, AFA, Hubley, R & Green P. RepeatMasker Open-4.0 [Internet]. 2010. Available from: www.repeatmasker.org

101. Smit, AFA, Hubley R. RepeatModeler Open-1.0 [Internet]. 2010. Available from: www.repeatmasker.org

102. Bao W, Kojima KK, Kohany O. Repbase Update, a database of repetitive elements in eukaryotic genomes. Mob DNA. 2015;6:11.

103. Kurtz S, Phillippy A, Delcher AL, Smoot M, Shumway M, Antonescu C, et al. Versatile and open software for comparing large genomes. Genome Biol. 2004;5:R12.

104. Nattestad M, Chin C-S, Schatz MC. Ribbon: Visualizing complex genome alignments and structural variation. bioRxiv. 2016;0344:82123.

